# Different Routes for Direct Interspecies Electron Transfer with Diverse Electron-Accepting Partners

**DOI:** 10.1101/2022.04.27.489562

**Authors:** Dawn E. Holmes, Jinjie Zhou, Jessica A. Smith, Caiqin Wang, Xinying Liu, Derek R. Lovley

## Abstract

Direct interspecies electron transfer (DIET) may be most important in methanogenic environments, but studies to date that have examined the mechanisms for electron transfer from electron-donating partners have primarily focused on co-cultures in which fumarate was the terminal electron acceptor. To better understand DIET with methanogens, the transcriptome of *Geobacter metallireducens* during DIET-based growth with *Geobacter sulfurreducens* reducing fumarate was compared with the transcriptome of *G. metallireducens* grown in co-culture with diverse *Methanosarcina* species. The transcriptome of *G. metallireducens* co-cultured with *G. sulfurreducens* was significantly different from those with *Methanosarcina*. Furthermore, the transcriptome of *G. metallireducens* grown with *Methanosarcina barkeri*, which lacks outer-surface cytochromes, was different from the transcriptome of *G. metallireducens* co-cultured with *Methanosarcina acetivorans* or *Methanosarcina subterranea*, which have an outer-surface *c*-type cytochrome that serves as an electrical connect for DIET. Differences in *G. metallireducens* expression patterns for *c*-type cytochrome genes were particularly notable. The impact of deleting *G. metallireducens c*-type cytochrome genes depended upon the electron-accepting partner. Multiple cytochromes that *G. metallireducens* required for DIET with *G. sulfurreducens* and for Fe(III) oxide reduction were not required for DIET with the *Methanosarcina* species. Porin-cytochrome complexes and e-pili were required for optimal DIET with all partners. *Shewanella oneidensis*, another electroactive microbe with abundant outer-surface *c*-type cytochromes, did not grow via DIET. The results demonstrate that the presence of outer-surface cytochromes does not necessarily confer the capacity for DIET, and also emphasize the substantial impact of the physiology of the electron-accepting partner on the physiology of the electron-donating DIET partner.

## Introduction

Direct interspecies electron transfer (DIET) is proposed to play a major role in the global cycling of methane, an important greenhouse gas, and in the generation of methane biofuel from organic wastes. Syntrophic communities producing or consuming methane in anaerobic soils and sediments share electrons via DIET (1, 2). Promoting DIET in anaerobic digesters through changes in digester design or with the addition of electrically conductive materials can accelerate anaerobic digestion (3, 4). Thus, understanding the mechanisms for electron exchange during DIET could contribute to better modeling of carbon and electron flux in natural anaerobic environments and could suggest better strategies for enhancing DIET during organic waste treatment.

Until now, studies on the routes for interspecies electron transfer during DIET have primarily focused on the mechanisms of electron uptake by the electron-accepting partner (5-9). These studies have demonstrated that the electrical connects on the outer cell surface of electron-accepting partners can be dramatically different. For example, *Geobacter sulfurreducens* displays a network of electrically conductive pili (e-pili) and outer surface multi-heme *c*-type cytochromes to facilitate electron uptake during DIET (5, 6, 10). In contrast, some methanogens can express electrically conductive archaella (11), but archaella are not known to participate in DIET and do not form a multi-fiber conductive extracellular mesh like *Geobacter* e-pili. Furthermore, some methanogens, such as *Methanothrix harundinacea, Methanobacterium electrotrophus* strain YSL, and *Methanosarcina barkeri*, are capable of DIET despite a lack of outer-surface *c*-type cytochromes (7, 8, 12-16). The electrical contacts for DIET in these methanogens are unknown. However, another methanogen, *Methanosarcina acetivorans*, expresses an outer-surface multi-heme *c*-type cytochrome, MmcA, that is a key electrical contact for DIET (9).

This major difference in mechanisms for DIET-based electron uptake between *M. barkeri* and *M. acetivorans* is just one of the physiological dissimilarities between two clades of *Methanosarcina* species. Type I *Methanosarcina*, like *M. barkeri*, predominate in waste digesters and other high-energy environments and have different mechanisms for energy conservation than Type II *Methanosarcina*, such as *M. acetivorans*, that are better adapted for growth in organic-poor soils and sediments (15). It might be expected that differences in the diversity of electrical contacts on electron-accepting partners for DIET could influence the routes for electron transfer from electron-donating partners.

Furthermore, the majority of studies on DIET mechanisms have focused on co-cultures in which *G. sulfurreducens* reducing fumarate was the electron-accepting partner. DIET to a fumarate-reducing partner is unlikely to be important in anaerobic environments because fumarate is not a commonly abundant electron acceptor. In addition, most simple organic substrates can be directly metabolized within individual fumarate-respiring microbes, which means that syntrophy is not required. In contrast, the inability of most methanogens to use multi-carbon electron donors other than acetate necessitates syntrophic metabolism with interspecies electron transfer. The electron carriers that provide electrons to support fumarate reduction during DIET could be much different than those required to promote the reduction of carbon dioxide to methane because the mid-point potential of the carbon dioxide/methane redox couple is much more negative than the fumarate/succinate couple.

A diversity of electron-donating partners for DIET have been identified, but in most instances genetically tractable isolates for mechanistic studies are not available. For example, pure cultures are not available for the archaea proposed to function as electron-donating partners for DIET in consortia catalyzing anaerobic methane oxidation coupled to sulfate reduction (17-19). Tools for genetic manipulation have not been developed for bacteria, such as *Syntrophus aciditrophicus* (20), *Rhodoferax ferrireducens* (8) and *Desulfovibrio* strain JY (21), shown to serve as the electron-donating partner for DIET in defined co-cultures.

Therefore, *Geobacter metallireducens*, which is genetically tractable (22), has served as the primary pure culture model for an electron-donating partner in DIET. *G. metallireducens* is an attractive model for DIET studies because of its environmental relevance. *Geobacter* species are important electron-donating DIET partners in some anaerobic waste digesters (4, 23) and in rice paddy soils, an important source of atmospheric methane (24). *G. metallireducens* is a good candidate for studying DIET in defined systems because it can oxidize ethanol and other short-chain alcohols with electron transfer to extracellular electron acceptors without generating H_2_ or formate (5, 6, 10, 25, 26).

Here we report that DIET transcriptomes suggest that the physiology of the electron-accepting partner directly impacts the physiology of *G. metallireducens* serving as the electron-donating partner. Gene deletion studies suggested different routes for electron transport, depending on which partner *G. metallireducens* was paired with. We also demonstrate that in many instances *G. metallireducens*/*Methanosarcina* co-cultures adapted over time to gene deletions that initially disrupted DIET. Attempts to simplify the study of DIET extracellular electron exchange by initiating DIET co-cultures with *Shewanella oneidensis*, an electroactive microbe with a less complicated array of outer-surface electron transport options, were unsuccessful.

## Results and Discussion

### Gene expression patterns suggest that the electron-accepting partner influences routes for DIET

DIET between *G. metallireducens* and *G. sulfurreducens* was compared to DIET between *G. metallireducens* and three *Methanosarcina* species: *M. barkeri* (MB), a Type I *Methanosarcina* that lacks outer-surface cytochromes (15) and *M. acetivorans* (MA), and *M. subterranea* (MS), Type II *Methanosarcina* with the outer-surface multi-heme *c*-type cytochrome MmcA that serves an essential electrical contact for DIET (9).

Multidimensional scaling analysis with the biological coefficient of variation method revealed that the transcriptomes of *G. metallireducens* grown in DIET-based co-culture with *Methanosarcina* species were significantly different from transcriptomes of *G. metallireducens* grown in co-culture with *G. sulfurreducens* (Figure 1, Supplementary Tables S1-S3, and Supplementary Figure S1). Furthermore, the transcriptomes of *G. metallireducens* grown with the Type I *Methanosarcina, M. barkeri*, were significantly different from the *G. metallireducens* transcriptomes grown with the Type II *Methanosarcina* species, *M. acetivorans* and *M. subterranea*. The transcriptomes of *G. metallireducens* grown with either of the Type II *Methanosarcina* were similar (Figure 1). These results indicate that properties of the electron-accepting DIET partner substantially influence the physiological status of *G. metallireducens* during growth via DIET. Differences in gene expression suggested substantial differences in many aspects of *G. metallireducens* physiology (Supplementary Figure S2). Here we focus on the expression of possible outer-surface electrical connects for DIET.

**Figure 1.**
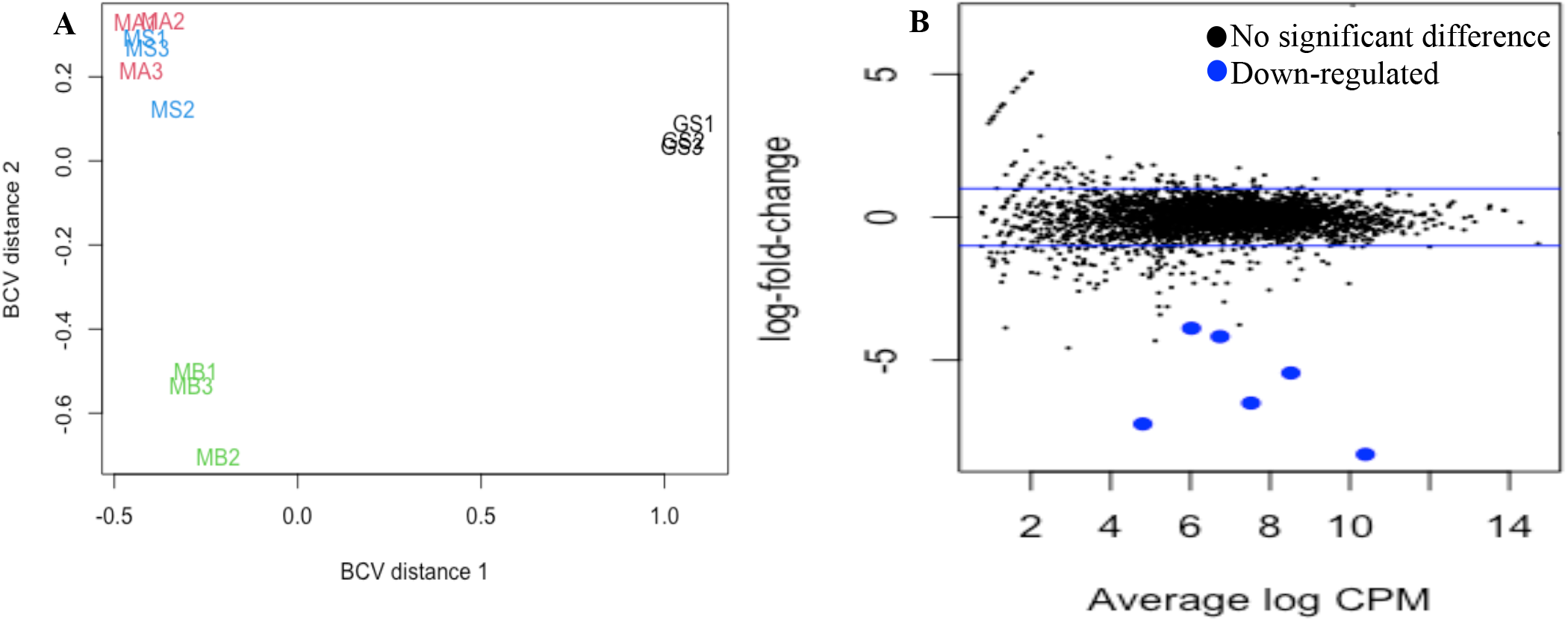
(A) Comparison of *G. metallireducens* RNAseq libraries from co-cultures with *Methanosarcina barkeri* (MB), *Methanosarcina acetivorans* (MA), *Methanosarcina subterranea* (MS), and *Geobacter sulfurreducens* (GS) using multidimensional scaling analysis with the biological coefficient of variation (BCV) method. (B) Mean-difference plots generated with plotMD from the LIMMA package in RStudio comparing *G. metallireducens* normalized transcripts from co-cultures grown with *M. acetivorans* or *M. subterranea* as the electron accepting partner. All libraries were done in triplicate and were normalized with the TMM (weighted trimmed means of M values) method.

Some of the most significant differences in *G. metallireducens* gene expression patterns during growth with the different types of electron-accepting partners were for genes coding for multi-heme *c*-type cytochromes (Figure 2, Tables 1-3, Supplementary Tables S1 and S2). The pattern of cytochrome gene expression was most similar in *G. metallireducens* grown with the two Type II *Methanosarcina* and the expression patterns between *G. metallireducens* and the Type I and Type II *Methanosarcina* were more similar to each other than to *G. metallireducens* grown with *G. sulfurreducens* as the electron-accepting partner. Therefore, similarities and differences in *G. metallireducens* cytochrome gene expression and the impact of targeted gene deletions on the ability of *G. metallireducens* to establish DIET with different electron-accepting partners were further evaluated.

**Table 1.** Differences in expression of genes coding for electron transfer proteins in *G. metallireducens* that were important for DIET when *G. sulfurreducens* was the electron-accepting partner. Values represent fold difference between co-cultures grown with *G. sulfurreducens* (GS) compared to co-cultures grown with the 3 different *Methanosarcina* species (*M. barkeri* (MB); *M. acetivorans* (MA); *M. subterranea* (MS)). p-values are available in Supplementary Table S2 but all comparisons shown have p-values that are <0.05. NS: no significant difference; Negative values show that the gene was more highly expressed by *G. metallireducens* grown in co-culture with the methanogen

| Locus ID | Annotation | Gene | Location | GS vs MB | GS vs MA | GS vs MS |
| --- | --- | --- | --- | --- | --- | --- |
| Gmet_0558 | c-type cytochrome protein, 27 hemes |  | Unknown | 18.28 | 19.68 | 13.50 |
| Gmet_0557 | c-type cytochrome protein, 4 hemes |  | Unknown | 18.02 | 22.42 | 24.19 |
| Gmet_2896 | c-type cytochrome protein, 4 hemes |  | Outer membrane/surface | 17.01 | 33.12 | 46.11 |
| Gmet_0930 | c-type cytochrome, 8 hemes |  | Outer membrane/surface | NS | -2.80 | -2.57 |
| Gmet_1868 | c-type cytochrome, 4 hemes |  | Outer membrane/surface | NS | 4.12 | 2.31 |
| Gmet_0571 | c-type cytochrome protein, 34 hemes |  | Outer membrane/surface | 12.63 | 3.29 | 2.72 |
| Gmet_0581 | c-type cytochrome protein, 34 hemes | *omcN* | Outer membrane/surface | 2.31 | NS | NS |
| Gmet_0911 | Porin protein from PccG complex | *ombG* | Outer membrane/surface | 80.67 | NS | 4.0 |
| Gmet_0912 | 8 heme periplasmic c-type cytochrome from PccG complex | *omaG* | Periplasmic | 96.91 | 2.01 | 3.58 |
| Gmet_0913 | 9 heme outer membrane c-type cytochrome from PccG | *omcG* | Outer membrane/surface | 70.53 | 2.34 | 3.43 |
| Gmet_0908 | Porin protein from PccF complex | *ombF* | Outer membrane/surface | -3.57 | -4.76 | -2.41 |
| Gmet_0909 | 9 heme periplasmic c-type cytochrome from PccF complex | *omaF* | Periplasmic | -6.50 | -7.08 | -3.81 |
| Gmet_0910 | 10 heme c-type cytochrome from PccF complex | *omcF* | Outer membrane/surface | -6.41 | -11.42 | -6.30 |
| Gmet_1399 | Type IV major pilin subunit, PilA | *pilA* | Outer membrane/surface | -4.31 | 1.64 | 1.76 |
| Gmet_1400 | Short pilin chaperone protein, Spc | *spc* | Outer membrane/surface | -7.85 | NS | NS |
| Gmet_2163 | Multicopper oxidase protein | *ompB* | Outer membrane/surface | 3.89 | 7.66 | 10.72 |
| Gmet_2928 | 7 heme c-type cytochrome protein from CbcABCDE | *cbcA* | Periplasmic | 11.09 | 4.34 | 4.13 |
| Gmet_2929 | b-type cytochrome from CbcABCDE complex | *cbcB* | Inner membrane | 7.85 | 6.08 | 5.22 |
| Gmet_2930 | 11 heme c-type cytochrome protein, CbcABCD complex | *cbcC* | Periplasmic | 2.16 | 2.0 | 2.14 |
| Gmet_2931 | Monoheme c-type cytochrome protein from CbcABCDE complex | *cbcD* | Periplasmic | 3.18 | 2.21 | 2.50 |
| **Gmet_2932** | Membrane protein from CbcABCDE complex | *cbcE* | Inner membrane | 4.46 | 1.74 | 1.80 |
| **Gmet_0252** | Monoheme *c*-type cytochrome protein | *coxB* | Inner membrane | 6.65 | 2.46 | 4.09 |
| **Gmet_0325** | 4 heme *c*-type cytochrome protein from CbcSTU-2 complex | *cbcS-2* | Periplasmic | 5.37 | 11.80 | 3.18 |
| **Gmet_0327** | *b*-type cytochrome from CbcSTU-1 complex | *cbcU-2* | Inner membrane | 2.25 | 9.48 | 4.64 |
| **Gmet_3519** | 4Fe-4S ferredoxin from CbcSTU-1 complex | *cbcT-1* | Periplasmic | NS | 3.47 | 3.44 |
| **Gmet_3520** | *b*-type cytochrome from CbcSTU-2 complex | *cbcU-1* | Inner membrane | 3.86 | 9.98 | 4.14 |
| **Gmet_0100** | Quinone oxidoreductase cytochrome, 9 hemes | *cbcL* | Inner membrane | 2.96 | 4.51 | 3.37 |
| **Gmet_3091** | *c*-type cytochrome protein, 2 hemes | *macA* | Periplasmic | 3.82 | NS | NS |
| **Gmet_3165** | *c*-type cytochrome protein, 3 hemes | *ppcC* | Periplasmic | 3.31 | 2.29 | NS |
| **Gmet_0580** | *c*-type cytochrome protein, 21 hemes |  | Outer membrane/surface | 3.02 | NS | NS |
| **Gmet_0600** | *c*-type cytochrome protein, 19 hemes |  | Outer membrane/surface | 2.21 | NS | NS |
| **Gmet_0575** | *c*-type cytochrome protein, 27 hemes |  | Outer membrane/surface | 2.01 | 3.03 | 2.86 |
| **Gmet_3088** | *c*-type cytochrome protein, 8 hemes |  | Unknown | 8.84 | 5.59 | 12.98 |
| **Gmet_2898** | *c*-type cytochrome protein, 8 hemes |  | Unknown | 6.81 | NS | NS |
| **Gmet_2470** | *c*-type cytochrome protein, 34 hemes |  | Unknown | 5.67 | 2.05 | 1.70 |
| **Gmet_2899** | *c*-type cytochrome protein, 9 hemes |  | Unknown | 5.43 | NS | NS |
| **Gmet_1087** | *c*-type cytochrome protein, 1 heme |  | Unknown | 4.46 | 8.40 | 6.56 |
| **Gmet_0601** | *c*-type cytochrome protein, 8 hemes |  | Unknown | 2.34 | NS | NS |
| **Gmet_0534** | *c*-type cytochrome protein, 5 hemes |  | Unknown | NS | 3.36 | 3.10 |
| **Gmet_0121** | *c*-type cytochrome protein, 2 hemes |  | Unknown | NS | 2.12 | 2.09 |

**Table 2.** Genes coding for electron transport proteins in *G. metallireducens* that had greater than 2 fold differences in expression in co-cultures grown with *M. barkeri* compared to growth with *G. sulfurreducens* (GS) and the Type II *Methanosarcina* (*M. acetivorans* (MA) and *M. subterranea* (MS)). p-values are available in Supplementary Table S2 but all comparisons shown have p-values that are <0.05. NS: no significant difference

| Locus ID | Annotation | Gene | Location | MB vs GS | MB vs MA | MB vs MS |
| --- | --- | --- | --- | --- | --- | --- |
| Gmet_2119 | Cytochrome *b* from PcmABCDEFGF complex | *pcmC* | Inner membrane | 104.99 | 9.93 | NS |
| Gmet_2117 | Membrane protein from PcmABCDEFGF complex | *pcmA* | Inner membrane | 10.83 | 9.17 | 7.19 |
| Gmet_1019 | *c*-type cytochrome, 2 hemes | *narC* | Inner membrane | 8.44 | NS | NS |
| Gmet_0330 | Monoheme *c*-type cytochrome | *narH* | Inner membrane | 5.37 | NS | NS |
| Gmet_1809 | *c*-type cytochrome, 5 hemes | *actA* | Inner membrane | 5.15 | NS | NS |
| Gmet_1923 | *b*-type cytochrome from CbcWXV complex | *cbcW* | Inner membrane | 3.84 | 1.54 | NS |
| Gmet_1922 | Rieske 2Fe-2S protein from CbcWXV complex | *cbcV* | Inner membrane | 1.92 | 1.64 | 2.13 |
| Gmet_2120 | Cytochrome *b* from PcmABCDEFGF complex | *pcmD* | Inner membrane | NS | 4.66 | NS |
| Gmet_2122 | 4 heme *c*-type cytochrome from PcmABCDEFGF complex | *pcmF* | Periplasmic | 123.26 | 6.85 | 83.58 |
| Gmet_2123 | 4 heme *c*-type cytochrome from PcmABCDEFGF complex | *pcmG* | Periplasmic | 8.94 | 20.04 | 47.60 |
| Gmet_3166 | *c*-type cytochrome, 2 hemes | *ppcB* | Periplasmic | 7.65 | 3.13 | 3.37 |
| Gmet_1210 | *c*-type cytochrome, 2 hemes | *ccpB* | Periplasmic | 7.59 | NS | 2.11 |
| Gmet_2156 | *c*-type cytochrome, 9 hemes |  | Periplasmic | 7.19 | 2.87 | NS |
| Gmet_1703 | *c*-type cytochrome, 7 hemes |  | Periplasmic | 6.71 | 7.63 | 3.59 |
| Gmet_0909 | 9 heme *c*-type cytochrome from PccF complex | *omaF* | Periplasmic | 6.50 | NS | NS |
| Gmet_1846 | *c*-type cytochrome, 2 hemes | *ppcE* | Periplasmic | 6.12 | 5.52 | 3.33 |
| Gmet_0328 | *c*-type cytochrome, 2 hemes | *narC* | Periplasmic | 5.80 | 2.62 | 2.50 |
| Gmet_0335 | *c*-type cytochrome, 3 hemes | *ppcF* | Periplasmic | 5.52 | 2.95 | 2.64 |
| Gmet_2902 | *c*-type cytochrome, 2 hemes | *ppcA* | Periplasmic | 4.31 | 2.42 | 3.16 |
| Gmet_1924 | 5 heme *c*-type cytochrome from CbcWXV complex | *cbcX* | Periplasmic | 4.24 | 2.98 | 2.85 |
| **Gmet_0155** | Monoheme *c*-type cytochrome |  | Periplasmic | 3.50 | NS | NS |
| **Gmet_0170** | *c*-type cytochrome, 10 hemes |  | Periplasmic | 3.12 | NS | NS |
| **Gmet_0142** | *c*-type cytochrome, 8 hemes |  | Periplasmic | 2.42 | NS | NS |
| **Gmet_0910** | 10 heme *c*-type cytochrome from PccF complex | *omcF* | Outer membrane/surface | 6.41 | NS | NS |
| **Gmet_0908** | Porin protein from PccF complex | *ombF* | Outer membrane/surface | 3.57 | NS | NS |
| **Gmet_1868** | *c*-type cytochrome, 4 hemes |  | Outer membrane/surface | NS | 4.41 | 2.35 |
| **Gmet_2121** | Rieske 2Fe-2S protein from PcmABCDEFGF complex | *pcmE* | Unknown | 53.94 | 11.03 | NS |
| **Gmet_2118** | 4Fe-4S ferredoxin protein from PcmABCDEFGF complex | *pcmB* | Unknown | 18.16 | 3.76 | NS |
| **Gmet_1647** | *c*-type cytochrome, 2 hemes |  | Unknown | 9.35 | NS | NS |
| **Gmet_1094** | *c*-type cytochrome, 2 hemes | *cccA* | Unknown | 6.02 | NS | NS |
| **Gmet_1197** | *c*-type cytochrome, 5 hemes |  | Unknown | 5.64 | NS | NS |
| **Gmet_1191** | Monoheme *c*-type cytochrome |  | Unknown | 4.49 | NS | NS |
| **Gmet_1814** | Monoheme *c*-type cytochrome | *actE* | Unknown | 4.16 | NS | NS |
| **Gmet_3506** | *c*-type cytochrome, 2 hemes |  | Unknown | 3.49 | NS | NS |
| **Gmet_2432** | Monoheme P460 *c*-type cytochrome |  | Unknown | 2.71 | NS | NS |
| **Gmet_2896** | *c*-type cytochrome, 4 hemes |  | Unknown | NS | 2.01 | 2.63 |

**Table 3.**
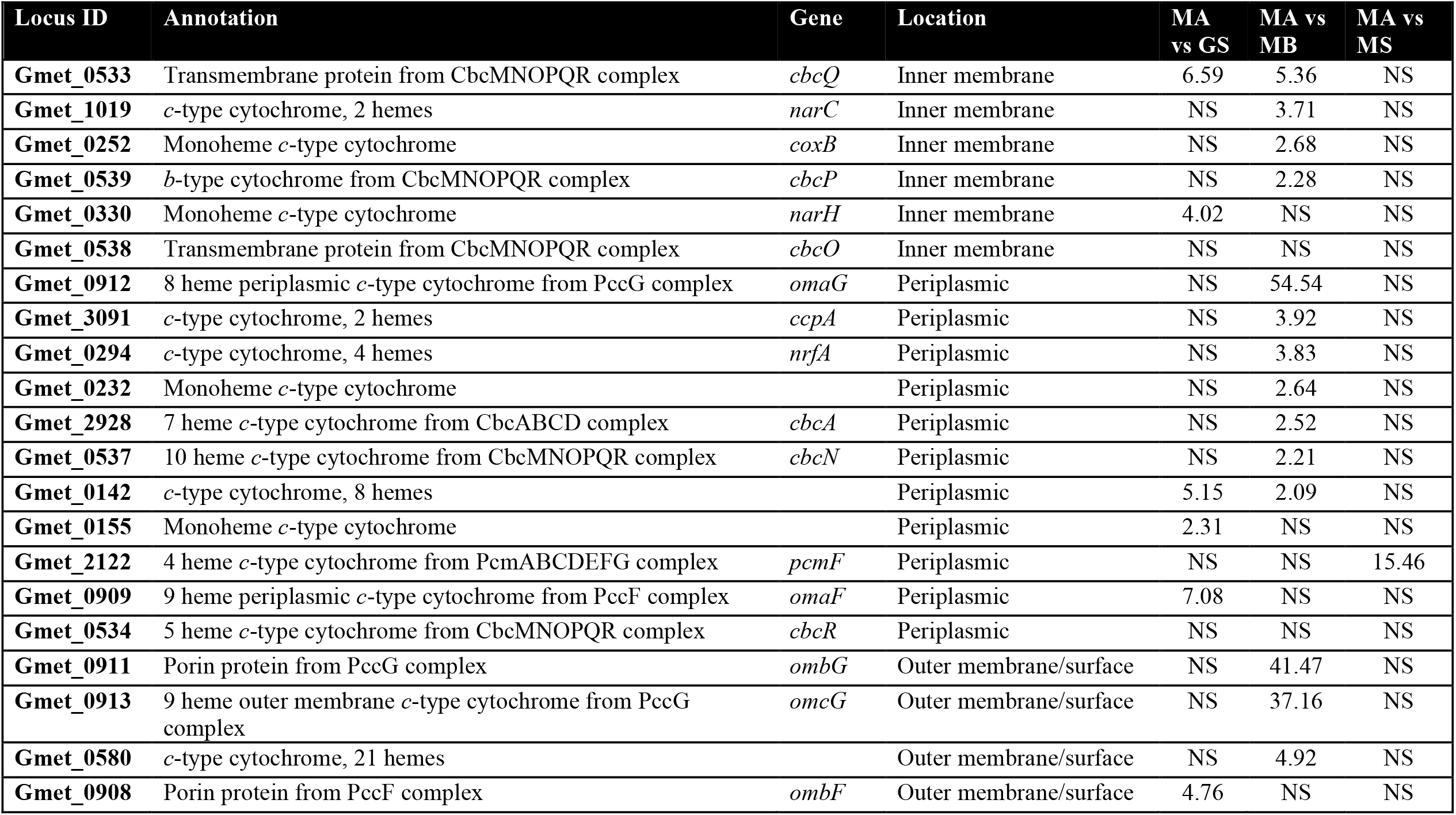
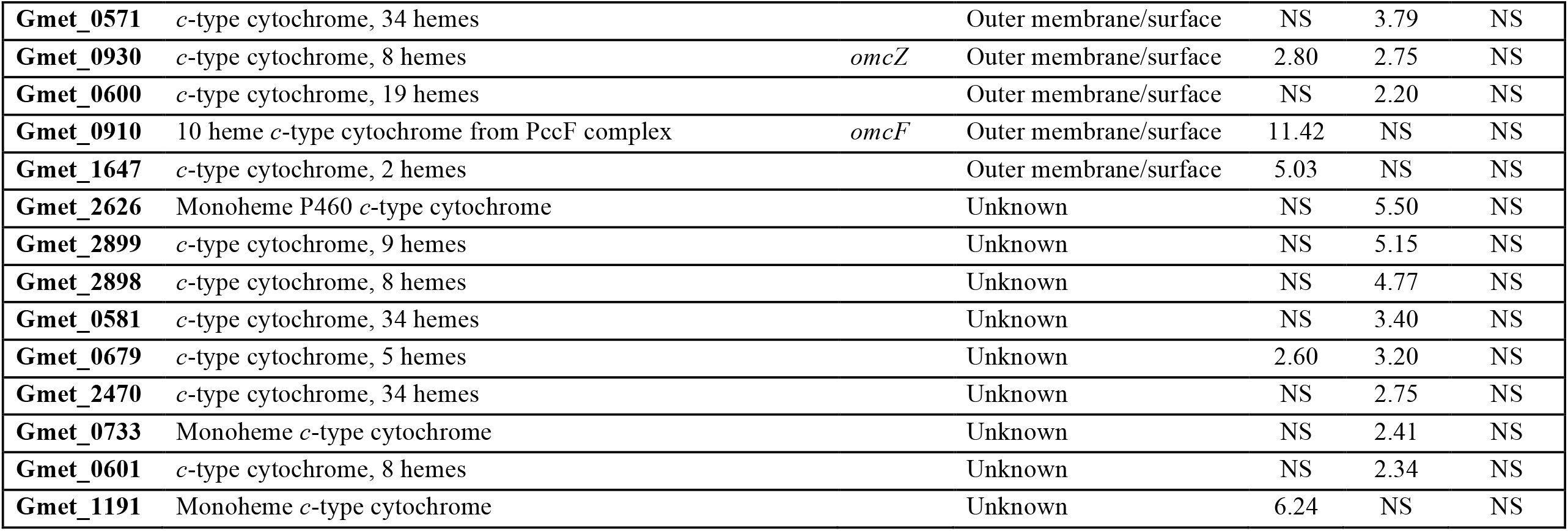
Genes coding for electron transport proteins in *G. metallireducens* that had greater than 2 fold differences in expression in co-cultures grown with the Type II *Methanosarcina* (*M. acetivorans* (MA)) than with *G. sulfurreducens* (GS), *M. barkeri* (MB), or *M. subterranea* (MS). p-values are available in Supplementary Table S2 but all comparisons shown have p-values that are <0.05. NS: no significant difference

**Figure 2.**
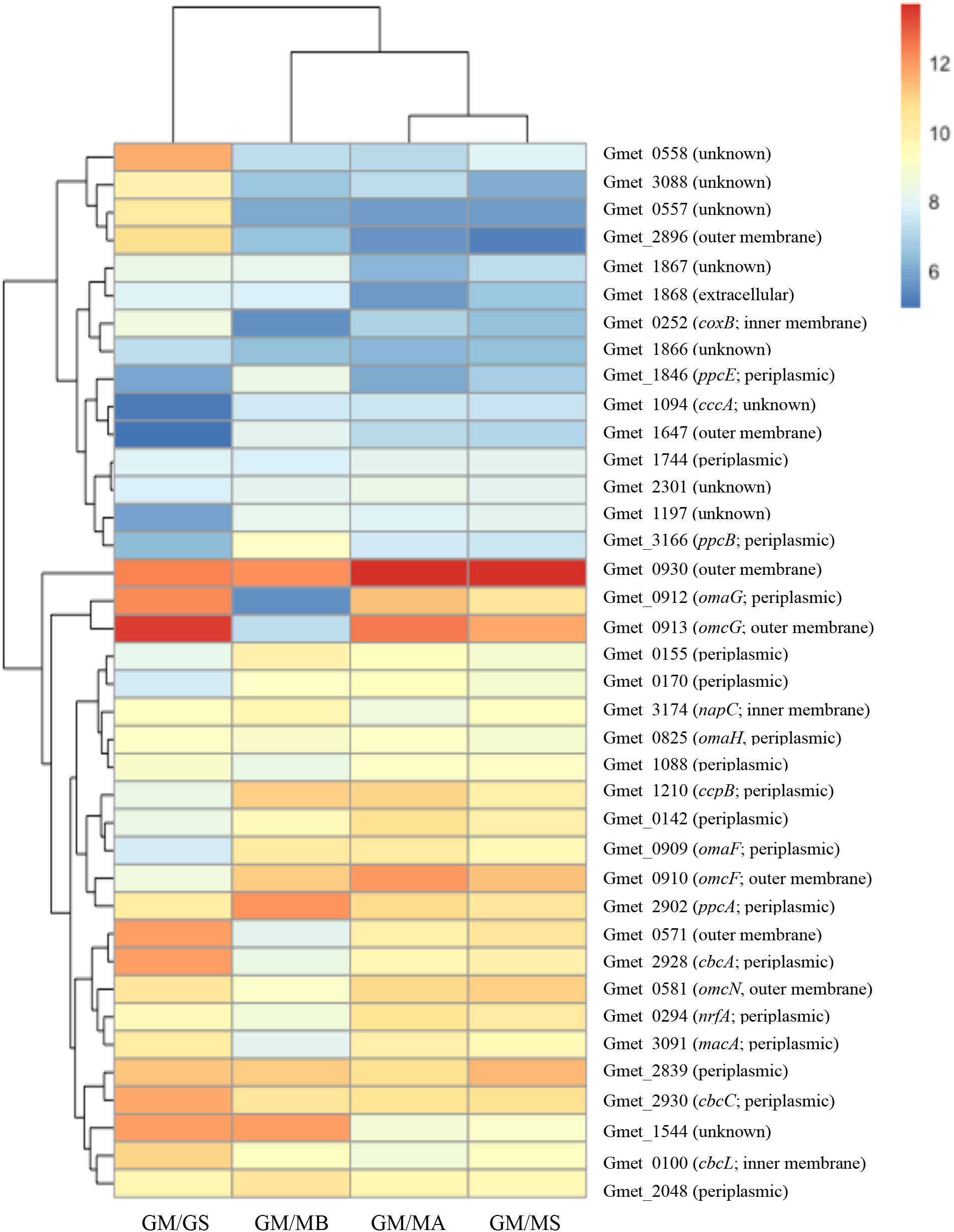
Log_2_ read values normalized with the TMM method for *G. metallireducens c*-type cytochrome genes from 4 different RNAseq co-culture libraries (GM/GS: *G. metallireducens* and *G. sulfurreducens*, GM/MB: *G. metallireducens* and *M. barkeri*, GM/MA: *G. metallireducens* and *M. acetivorans*, GM/MS: *G. metallireducens* and *M. subterranea*) that had log_2_ values above the median TMM values.

### Outer surface *c*-type cytochromes

Multi-heme, *c*-type cytochromes positioned on the outer cell surface can serve as a key electrical contact for interspecies electron transfer (27). Gene expression patterns and the differential impact of deleting genes for outer-surface *c*-type cytochromes suggested that the relative importance of individual *G. metallireducens* outer-surface cytochromes depended upon which microbe served as the electron-accepting partner for DIET.

For example, a major difference between *G. sulfurreducens* or *Methanosarcina* species serving as the electron-accepting partner was the apparent role of the multi-heme *c*-type cytochromes Gmet_0558, Gmet_0557, and Gmet_2896. Software designed to predict cellular localization did not specify a location for these cytochromes. However, the presence of a transmembrane helix in Gmet_0557 suggests that it is membrane associated, and Gmet_2896 cytochrome was recovered in outer membrane preparations of *G. metallireducens* (28, 29). The large number of hemes (27) predicted for Gmet_0558 and the fact that it has high homology (67% amino acid identity) to OmcO, an outer membrane *c*-type cytochrome in *G. sulfurreducens* (30), also suggests that it may be an outer-surface electrical contact.

The genes for all three of these cytochromes were more highly expressed when *G. sulfurreducens* was the electron-accepting partner than with any of the *Methanosarcina* species (Table 1). Deleting any of these three genes had no impact on establishing co-cultures with the three *Methanosarcina* species (Supplementary Figure S3). However, co-cultures initiated with *G. sulfurreducens* and Gmet_0557- or Gmet_2896-deficient strains did not grow and co-cultures initiated with a Gmet_0558-deficient mutant had a more extensive lag period than co-cultures initiated with wild-type *G. metallireducens* (Figure 3A). Even after further adaption with three more transfers, the co-cultures with the Gmet_0558-deficient strain reduced fumarate at rates that were 2.3 times slower (p=1.15×10^−5^) than co-cultures with wild-type *G. metallireducens* (Figure 3B). These results suggest that Gmet_0558, Gmet_0557, and Gmet_2896 are all important for extracellular electron transfer when *G. sulfurreducens* is the electron-accepting partner, but they are not essential for DIET with *Methanosarcina* with or without outer-surface *c*-type cytochrome electrical connects. Previous gene-deletion studies indicated that Gmet_0558, Gmet_0557, and Gmet_2896 are required for growth with insoluble Fe(III) oxide as the electron acceptor (28). Thus, the concept that *Geobacter* extracellular electron transfer to other microbial species requires the same outer-surface electron transport components as electron transport to Fe(III) oxide does not appear to hold when *Methanosarcina* are the partner for DIET.

**Figure 3.**
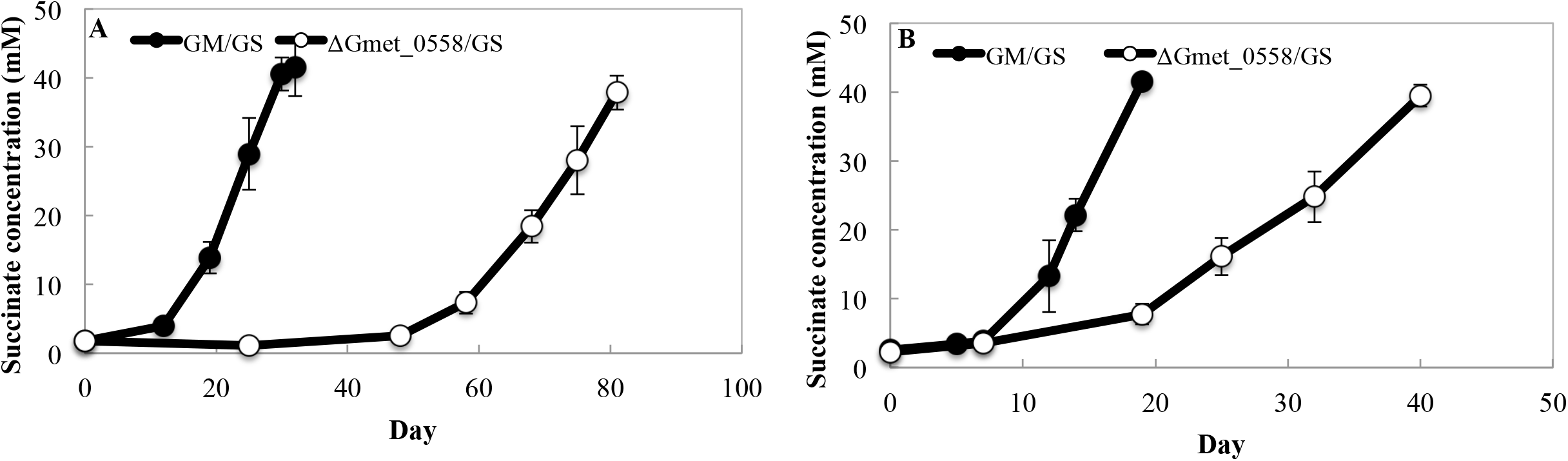
Succinate production by co-cultures grown with wild-type *G. metallireducens*, or the deletion mutant strain ΔGmet_0558, and *G. sulfurreducens* during (A) the first transfer when aggregates first became established or (B) the fourth transfer after aggregates had been established. Ethanol (10 mM) was provided as the electron donor and fumarate (45 mM) was provided as the electron acceptor. Error bars represent the standard deviation from triplicate cultures.

Gmet_0930 represents another instance in which an outer-surface *c*-type cytochrome was essential for DIET to *G. sulfurreducens* and Fe(III) oxide reduction, but not for DIET with *Methanosarcina* species. Gmet_0930, encodes an 8-heme outer-surface *c*-type cytochrome that is homologous to OmcZ (GSU2076) in *G. sulfurreducens*; they have 45% amino acid identity, they both have 8 hemes, and they both are predicted to be extracellular and have putative immunoglobulin like fold domains. Deletion of the gene for OmcZ had no impact on Fe(III) oxide reduction in *G. sulfurreducens* (31), but *G. metallireducens* requires Gmet_0930 to reduce Fe(III) oxide (28).

Gmet_0930 was one of the most highly expressed *G. metallireducens c*-type cytochrome genes with all electron-accepting partners (Figure 2, Supplementary Table S1). Co-cultures could not be established with *G. sulfurreducens* and a strain of *G. metallireducens* in which Gmet_0930 was deleted. Initiating co-cultures with the Gmet_0930-deficient strain delayed methane production by all three of the *Methanosarcina* strains (Figure 4). Although Gmet_0930 was not as essential for DIET with *Methanosarcina* species as it was for DIET with *G. sulfurreducens*, it did appear to optimize DIET because even after 4 transfers the *Methanosarcina* co-cultures initiated with the Gmet_0930-deficient strain grew 1.5-2-fold slower than co-cultures initiated with wild-type *G. metallireducens* (Supplementary Figure S4). The finding that deletion of Gmet_0930 inhibits DIET with *G. sulfurreducens*, as well as Fe(III) oxide reduction (28), further supports the conclusion from the study of Gmet_0558, Gmet_0557, and Gmet_2896 that there are differences between *G. metallireducens* electron transport to Fe(III) oxide and *G. sulfurreducens* compared to electron transport to *Methanosarcina* species.

**Figure 4.**
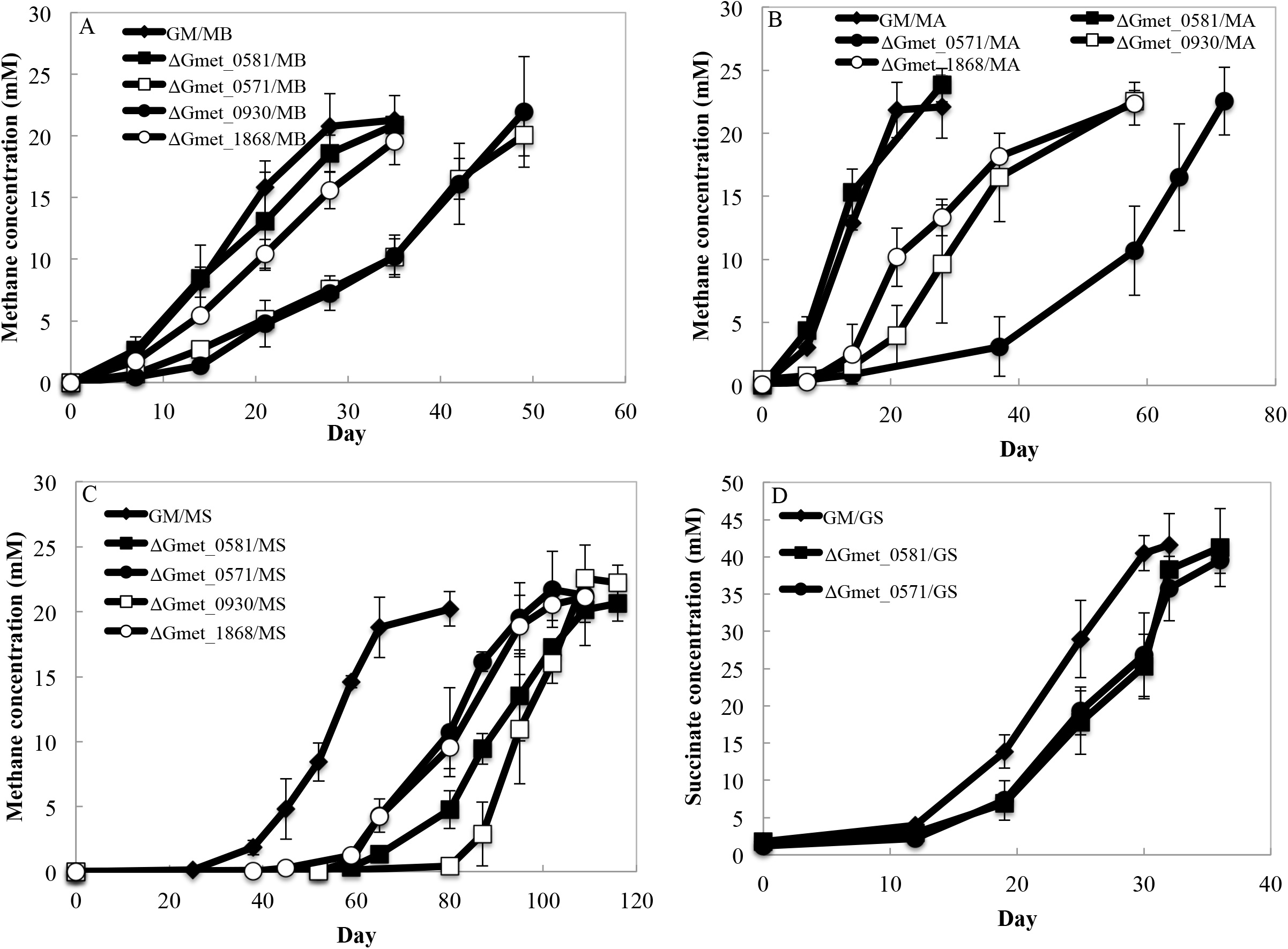
Methane or succinate production during the first transfer by co-cultures grown with various species of *Methanosarcina* or *G. sulfurreducens* and *G. metallireducens* strains lacking the *c*-type cytochromes encoded by Gmet_0571, Gmet_0581, Gmet_1868, or Gmet_0930. (A) Co-cultures with *M. barkeri*; (B) Co-cultures with *M. acetivorans*; (C) Co-cultures with *M. subterranea*; (D) Co-cultures with *G. sulfurreducens*. Error bars represent the standard deviation from triplicate cultures.

In a similar manner, the four-heme outer-surface *c*-type cytochrome Gmet_1868, which *G. metallireducens* requires for Fe(III) oxide reduction (28), appeared to be essential for DIET with *G. sulfurreducens*, but only helpful for DIET with the *Methanosarcina* species. Transcript abundance for Gmet_1868 was more than 2-fold higher than the median normalized TMM (weighted trimmed means of M values) values for the *G. metallireducens*/*G. sulfurreducens* and *G. metallireducens*/*M. barkeri* co-cultures, but the number of Gmet_1868 transcripts in both Type II *Methanosarcina* co-cultures was slightly below the TMM median log_2_ value (Supplementary Table S1). Co-cultures initiated with a Gmet_1868-deficient strain and *G. sulfurreducens* did not grow. The lag phase for the Gmet_1868-deficient strain and *Methanosarcina* co-cultures was longer than the wild-type co-cultures (Figure 4) and even after 4 transfers, the co-cultures initiated with the Gmet_1868-deficient strain and the *Methanosarcina* species grew at rates that were 1.4-3.0 fold slower than the wild-type co-culture (Supplementary Figure S4).

The 34-heme outer-surface *c*-type cytochrome Gmet_0571, which was highly expressed during growth with all electron-accepting partners (Figure 2) appeared to be more important for DIET with the *Methanosarcina* species than *G. sulfurreducens*. Co-cultures initiated with *G. sulfurreducens* and a strain of *G. metallireducens* in which the Gmet_0571 gene was deleted grew similarly to wild-type co-cultures (Figure 4 and Supplementary Figure S4). In contrast, co-cultures with the Gmet_0571-deficient strain and the *Methanosarcina* were substantially delayed in methane production compared to co-cultures initiated with wild-type *G. metallireducens* (Figure 4). With subsequent transfer, the ΔGmet_0571/MB, ΔGmet_0571/MA and ΔGmet_0571/MS co-cultures adapted somewhat to the loss of Gmet_0571, but they continued to grow 1.6-3.0 times slower than the co-cultures established with wild-type *G. metallireducens* (Supplementary Figure S4). These results suggest that Gmet_0571 is required for optimal DIET with *Methanosarcina* species, but is not necessary for DIET with *G. sulfurreducens* or for growth with Fe(III) oxide provided as the electron acceptor (28).

A different pattern of gene deletion phenotypes was observed with Gmet_0581, another putative 34-heme *c*-type cytochrome likely to be positioned on the outer cell surface. Gmet_0581 shares 63% amino acid identity with the outer membrane *c*-type cytochrome OmcN of *G. sulfurreducens* and has a surface-associated immunoglobulin-like domain. It was highly expressed with all electron-accepting partners. Deletion of Gmet_0581 impacted co-culture establishment with the Type II *Methanosarcina* but not with *M. barkeri* or *G. sulfurreducens* (Figure 4). However, after 4 transfers, co-cultures with the Type II *Methanosarcina* and the Gmet_0581-deficient strain grew at rates that were comparable to co-cultures established with wild-type *G. metallireducens* (Supplementary Figure S4).

### Porin-cytochrome complexes

Transcriptomic and gene deletion studies of the porin-cytochrome (Pcc) complexes also suggested that there are different requirements for DIET to *Methanosarcina* species versus DIET to *G. sulfurreducens. G. metallireducens* has genes for three Pcc complexes: PccF (Gmet_0908-0910), PccG (Gmet_0911-0913), and PccH (Gmet_0825-0827). Pcc complexes are thought to be necessary for electron transport across the outer membrane of *Geobacter* species (27, 32). The full suite of genes coding for components of the PccH complex was not highly expressed by *G. metallireducens* in any of the co-culture conditions, indicating that PccH was not important for DIET. However, some of the most substantial differences in *G. metallireducens* gene transcript abundance in co-cultures with *G. sulfurreducens* versus *M. barkeri* as the electron-accepting partner were for the genes that code for components of the PccG porin-cytochrome complex (Table 1). The genes from the PccF complex, on the other hand, were more highly expressed by *G. metallireducens* grown in the three *Methanosarcina* co-cultures (Table 1).

Initiating co-cultures with strains of *G. metallireducens* in which the gene for OmcF or OmcG, the outer-membrane cytochromes of the PccF and PccG conduits, was deleted delayed the initiation of methane production in co-cultures established with each of the *Methanosarcina* species (Figure 5A-C). There was also a slight lag in methane production in co-cultures established with *G. sulfurreducens* and the OmcF*-* and OmcG*-* deficient strains of *G. metallireducens* (Figure 5D). Deletion of the gene coding for the periplasmic-facing *c*-type cytochrome, *omaG* (Gmet_0912) also delayed methane production with all three *Methanosarcina* species (Figure 5A-C), but did not have an impact on co-culture growth with *G. sulfurreducens* (Figure 5D).

**Figure 5.**
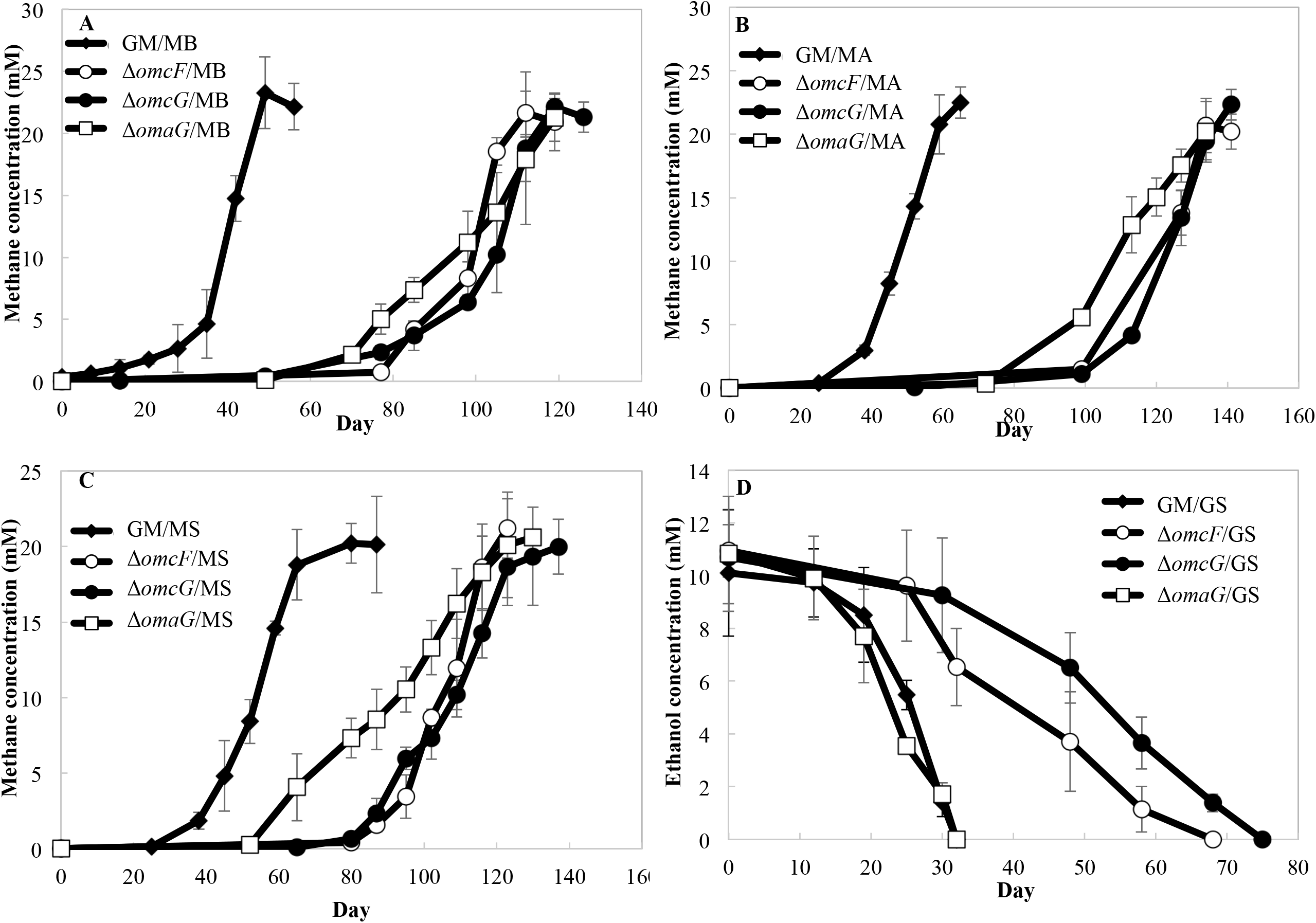
Methane production or ethanol consumption during the first transfer by co-cultures grown with various species of *Methanosarcina* or *G. sulfurreducens* and *G. metallireducens* strains lacking the porin associated outer membrane *c*-type cytochromes OmcF (Gmet_0910), OmcG (Gmet_0913) or the periplasmic porin associated cytochrome OmaG (Gmet_0912). (A) Co-cultures with *M. barkeri*; (B) Co-cultures with *M. acetivorans*; (C) Co-cultures with *M. subterranea*; (D) Co-cultures with *G. sulfurreducens*. Error bars represent the standard deviation from triplicate cultures.

After the co-cultures were passed through three additional sequential transfers, the co-cultures of *G. metallireducens* strain Δ*omcF* and the *Methanosarcina* species, strain Δ*omcF*/*G. sulfurreducens*, and strain Δ*omcG*/*G. sulfurreducens*, adapted to grow at rates that were similar to the wild-type (Supplementary Figure S5). However, even after this long-term adaption through multiple transfers, the co-cultures that had been initiated with *G. metallireducens* strain Δ*omcG* or strain Δ*omaG* and the *Methanosarcina* species grew ca. 2-4-fold slower than co-cultures initiated with wild-type *G. metallireducens* (Supplementary Figure S5). These results suggest that the PccG porin-cytochrome conduit has features that are optimized for DIET with *Methanosarcina* species, but that either porin-cytochrome conduit is suitable for DIET with *G. sulfurreducens*.

### e-Pili

The *G. metallireducens* gene for PilA, the pilin monomer that is assembled into e-pili (33), and the adjacent gene for Spc, a putative chaperone protein that facilitates pilus assembly (34), as well as many of the pilin accessory proteins (35), were highly expressed in *G. metallireducens* growing with each of the electron-accepting partners evaluated (Table 1 and Supplementary Table S4). Constructing *Geobacter* strains that express poorly conductive pili is a method for evaluating the evaluating the importance of pili conductivity in extracellular electron exchange without disrupting the proper expression of outer surface cytochromes that also play a key role in DIET (36-40). A strain of *G. metallireducens* that expresses poorly conductive pili was previously found to be defective in establishing DIET co-cultures with *G. sulfurreducens* (41) and the Type II *Methanosarcina* species, *M. acetivorans* (9). A co-culture initiated with *G. metallireducens* expressing poorly conductive pili and the Type I *Methanosarcina* species, *M. barkeri*, was delayed in converting ethanol to methane, but adapted to grow in co-culture at rates that were similar to the wild-type co-cultures after four transfers (Figure 6). These results suggest that e-pili are helpful, but not essential for establishing DIET with *M. barkeri*. Eliminating *G. metallireducens* e-pili expression had no impact on DIET between *G. metallireducens* and *Methanobacterium electrotrophus* strain YSL, which like *M. barkeri* lacks outer-surface *c*-type cytochromes (12). Thus, differences in outer-surface electron-accepting proteins between species that rely on outer-surface cytochromes (i.e. *G. sulfurreducens, M. acetivorans*) and species that do not (i.e. *M. barkeri, Methanobacterium electrotrophus* strain YSL) may influence whether e-pili are beneficial for establishing electrical contact.

**Figure 6.**
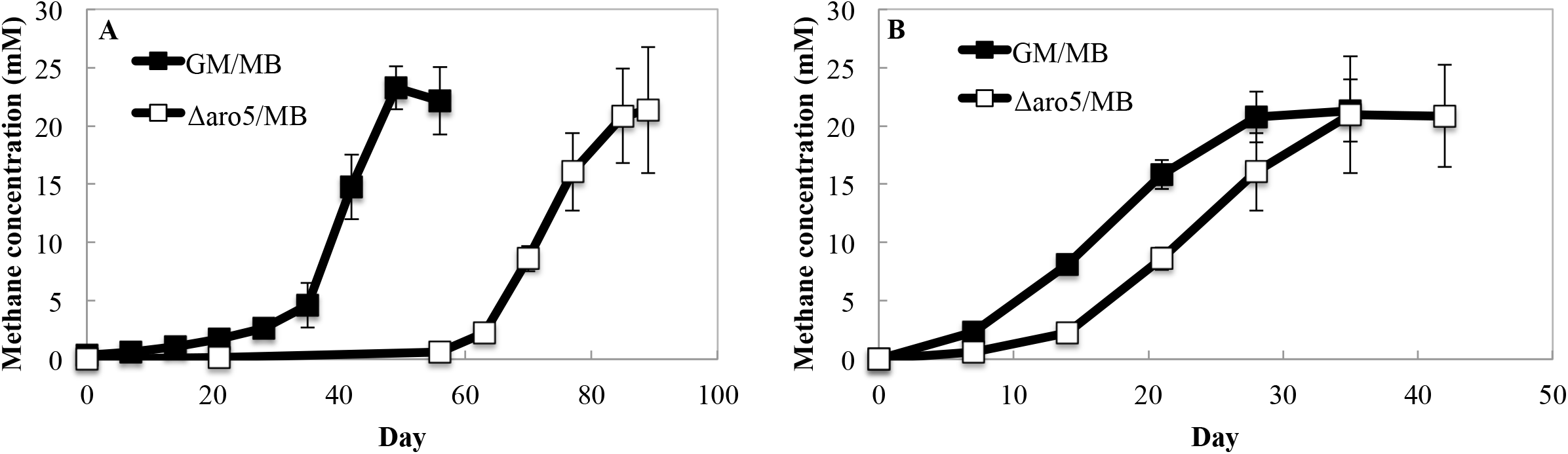
Methane production during growth of co-cultures with the *G. metallireducens* wild-type strain or the *G. metallireducens* Aro-5 mutant strain and *M. barkeri* during (A) the first transfer when aggregates became established or (B) the fourth transfer after aggregates had been established. Ethanol (20 mM) was provided as the electron donor. Error bars represent the standard deviation from triplicate cultures.

There were also substantial differences in expression of *G. metallireducens* genes coding for additional surface-associated proteins, with different electron-accepting partners (Supplementary Table S2). In particular, Gmet_2163, coding for a multi-copper protein homologous to OmpB, which *G. sulfurreducens* requires for Fe(III) oxide reduction (42), was more highly expressed in *G. metallireducens/G. sulfurreducens* co-cultures than it was in *G. metallireducens* co-cultures with the *Methanosarcina* species (Table 1). Further evaluation of the function of the multi-copper protein in DIET awaits construction of the appropriate mutant.

### Failure of *Shewanella oneidensis* to participate in DIET

In addition to illustrating likely differences between routes for DIET from *G. metallireducens* to *G. sulfurreducens* versus *Methanosarcina* species, the results above demonstrate that elucidating the routes for DIET with *G. metallireducens* is complex because of the high abundance of redox-active proteins on the outer surface, providing opportunities for adapting alternative routes for extracellular electron transfer. A simpler, yet still genetically tractable system to study DIET is desirable (27). Therefore, the possibility for *S. oneidensis* to serve as the electron-donating partner in DIET was investigated. Co-cultures were initiated with lactate as the electron donor and *S. oneidensis* as the electron-donating partner and either *M. barkeri, M. acetivorans, M. horonobensis*, or *M. subterranea* as the electron-accepting partner. None of the co-cultures grew or produced methane. Effective extracellular electron transfer by *S. oneidensis* is limited to soluble electron acceptors such as chelated Fe(III), metal ions, and electron shuttles, such as flavins (27, 43, 44). *S. oneidensis* lacks e-pili and primarily reduces extracellular particulate electron acceptors, such as Fe(III) oxides and electrodes, with flavins as an electron shuttle (45-47). Although soluble electron shuttles have been shown to facilitate interspecies electron transfer between *Geobacter* species with fumarate serving as the electron acceptor (48), the redox potential of reduced flavins may be too positive to support the reduction of carbon dioxide by methanogens. The apparent inability of *S. oneidensis* to participate in DIET with methanogens is consistent with the finding that *Shewanella* species are typically not reported to be abundant members of methanogenic communities.

### Implications

The results demonstrate that there may be substantial differences in the electron transport routes for DIET between *G. metallireducens* and *G. sulfurreducens* versus DIET between *G. metallireducens* and methanogens. Possible reasons for this include variations in key outer-surface electrical contacts and the difference in the redox potential required for the reduction of fumarate versus carbon dioxide. Differences in *G. metallireducens* transcriptomes and the impact of gene deletions on co-culture growth suggested that dissimilarities in electron uptake mechanisms between Type I and Type II *Methanosarcina* might also influence *G. metallireducens* physiology during DIET. The results also indicate that the common assumption that extracellular electron transfer to other cells goes through the same pathways as electron transfer to Fe(III) oxide is incorrect, especially for DIET to *Methanosarcina* species. A number of gene deletions previously shown to inhibit *G. metallireducens* growth on Fe(III) oxide had no impact on DIET-based growth with *Methanosarcina*.

These considerations highlight the need to develop models for DIET with microbes that are relevant to the environment of interest. Although *G. metallireducens/G. sulfurreducens* co-cultures with fumarate as the electron acceptor were convenient to develop the concept of DIET (5, 6, 10, 26, 49, 50), DIET is most likely to be environmentally significant under methanogenic conditions. Evaluation of *G. metallireducens* growing via DIET with other methanogens, such as *Methanobacterium* (12) and *Methanothrix* (13) species, that are physiologically and phylogenetically distinct from *Methanosarcina* species are required to more fully understand potential electron transport pathways for DIET. Further analysis of differences in gene expression patterns in diverse electron-donating microbes and electron-accepting methanogens will help identify the important electron transport pathways for DIET in methanogenic environments.

The finding that *S. oneidensis* was not capable of DIET with *Methanosarcina* species further demonstrates the often-ignored substantial differences in the extracellular electron transport capabilities of *Shewanella* and *Geobacter* species. Although *S. oneidensis* is commonly regarded as a model microbe equivalent to *Geobacter* species for the study of anaerobic biogeochemical processes and bioremediation, it lacks many of the metabolic capabilities of *Geobacter* species (27, 51, 52). More broadly, these results have implications for the inference for DIET based on metagenomic analyses, because the results suggest that the mere presence of genes for proteins, such as multi-heme *c*-type cytochromes, known to be key components for some forms of extracellular electron transfer, is not necessarily sufficient to confer the capacity for DIET.

## Materials and Methods

### Culture media and growth conditions

*Methanosarcina acetivorans* strain WWM1 (Δ*hpt*) (53) was routinely cultured under strict anaerobic conditions at 37°C in MA medium as previously described (9). *M. barkeri* MS (DSM 800), *M. horonobensis* HB-1 (DSM 21571) and *M. subterranea* DH-2 (15) were cultivated in the same medium with 1 g/L NaCl as opposed to the 4 g/L NaCl ordinarily found in MA medium. All *Methanosarcina* cultures were incubated in an N_2_-CO_2_ atmosphere (80:20, vol/vol) with acetate (40 mM) and methanol (20 mM) provided as substrates for growth.

*Geobacter metallireducens* GS-15 (ATCC 53774), *G. sulfurreducens* PCA (ATCC 51573), and *Shewanella oneidensis* MR-1 (ATCC 700550) were routinely cultured at 30°C under anaerobic conditions (N_2_:CO_2_, 80:20, vol/vol). Eighteen different *G. metallireducens c*-type cytochrome deletion mutant strains constructed as previously described (28) were obtained from our laboratory culture collection: ΔGmet_2896, ΔGmet_0558, ΔGmet_0534, ΔGmet_2928, ΔGmet_2930, ΔGmet_0557, ΔGmet_1868, ΔGmet_0930, ΔGmet_0679, ΔGmet_0825, ΔGmet_0913, ΔGmet_0910, ΔGmet_0912, ΔGmet_0571, ΔGmet_0232, Δ*fliC*, ΔGmet_2029 and ΔGmet_0581. All *G. metallireducens* strains were grown in freshwater medium (54) with ethanol (20 mM) provided as the electron donor and Fe(III) citrate (56 mM) provided as the electron acceptor. For growth of *G. sulfurreducens*, acetate (20 mM) was the donor and fumarate (40 mM) was the electron acceptor. *S. oneidensis* was grown with lactate (20 mM) provided as the electron donor and Fe(III) citrate (56 mM) provided as the electron acceptor. Growth of *Shewanella* required supplementation of 22 mg/L L-arginine, 22 mg/L L-glutamine, and 44 g/L DL-serine to the medium.

For co-culture experiments with *Methanosarcina* species, *G. metallireducens* strains and either *M. acetivorans, M. barkeri*, or *M. subterranea* were grown with 20 mM ethanol provided as the electron donor and carbon dioxide as the electron acceptor at 30°C as previously described (9, 14, 15). Attempted co-cultures with *S. oneidensis* and the various *Methanosarcina* were grown under the same conditions but lactate (20 mM) was provided as the electron donor and the medium was supplemented with the three amino acids mentioned above. For co-culture experiments with *Geobacter* species, *G. metallireducens* strains and *G. sulfurreducens* were grown with ethanol (20 mM) provided as the electron donor and fumarate (45 mM) or Fe(III) citrate (56 mM) as the electron acceptor in freshwater medium as previously described (5).

### Analytical techniques

A gas chromatograph equipped with a headspace sampler and a flame ionization detector (Clarus 600; PerkinElmer Inc., CA) was used to monitor ethanol concentrations. Methane in the headspace was measured by gas chromatography with a flame ionization detector (SHIMADZU, GC-8A) as previously described (55), and acetate concentrations were measured with a SHIMADZU high performance liquid chromatograph (HPLC) with an Aminex™ HPX-87H Ion Exclusion column (Bio-Rad, 300 mm × 7.8 mm) and an eluent of 8.0 mM sulfuric acid.

### Microscopy

Cells were routinely examined by phase-contrast and fluorescence microscopy (BV-2A filter set) with a Nikon E600 microscope.

### RNA Extraction

Cells were harvested from triplicate 50 mL cultures of *G. metallireducens* grown via DIET with *M. barkeri, M. acetivorans*, or *M. subterranea* during mid-exponential phase when ∼18 mM methane was detected in the headspace.

Cells were split into 50 mL conical tubes (BD Sciences), mixed with RNA Protect (Qiagen) in a 1:1 ratio, and pelleted by centrifugation at 3,000 x *g* for 15 minutes at 4°C. Pellets were then immediately frozen in liquid nitrogen and stored at -80°C. Total RNA was extracted from all samples as previously described (56), and mRNA was further enriched from all samples with the MICROB*Express* kit (Ambion).

### Illumina sequencing and data analysis

Directional multiplex libraries were constructed with RNA extracted from *G. metallireducens/M. barkeri* DIET co-cultures (GM/MB), *G. metallireducens/M. acetivorans* DIET co-cultures (GM/MA), and *G. metallireducens/M. subterranea* DIET co-cultures (GM/MS). The ScriptSeq™ v2 RNA-Seq Library Preparation Kit (Epicentre) was used to make the libraries according to the manufacturer’s instructions, and paired-end sequencing was performed on a Hi-Seq 2000 platform at the Deep Sequencing Core Facility at the University of Massachusetts Medical School in Worchester, Massachusetts.

Raw data was quality checked with FASTQC (http://www.bioinformatics.babraham.ac.uk/projects/fastqc/). All read information from libraries constructed for this study (GM/MB, GM/MA, GM/MS, GM/GS) is provided in Supplementary Material (Supplementary Table S5). Trimmomatic (57) was used to trim and filter all raw paired-end reads that were then merged with FLASH (58). Ribosomal RNA (rRNA) reads were removed from the libraries with SortMeRNA (59).

### Mapping of mRNA reads

Trimmed and filtered mRNA reads from the triplicate samples for the various co-culture conditions were mapped against the *G. metallireducens* GS-15 (NC_007517) genome downloaded from IMG/MER (img.jgi.doe.gov) using ArrayStar software (DNAStar). Reads were normalized and processed for differential expression studies using the edgeR package in Bioconductor (60).

## Data Availability

Illumina sequence reads have been submitted to the SRA NCBI database under BioProjects PRJNA727272, PRJNA722959, PRJNA828193, and PRJNA828235 and Biosamples SAMN19011638, SAMN18796025, SAMN27646189, and SAMN27646279.

## Acknowledgments

This research was supported by the Army Research Office and was accomplished under Grant Number W911NF-17-1-0345. The views and conclusions contained in this document are those of the authors and should not be interpreted as representing the official policies, either expressed or implied, of the Army Research Office or the U.S. Government.

## Contributions

Dawn Holmes and Jinjie Zhou conducted experiments, organized the study, and wrote the manuscript. Jessica Smith, Caiqin Wang, and Xinying Liu conducted experiments. Derek Lovley helped to organize the study and to write the manuscript.

## Ethics declarations

The authors do not declare any conflicts of interest.

