## Supplementary Text for "Different Routes for Direct Interspecies Electron Transfer with Diverse Electron-Accepting Partners"

In addition to apparent differences in extracellular electron transfer mechanisms utilized by *G. metallireducens* for growth via DIET with different electron accepting partners, a number of other genes involved in *Geobacter* physiology were also differentially expressed. Some of these differences are described here in Supplementary Material.

**Differential expression of quinone oxidoreductase complexes**

Studies of *G. sulfurreducens* have shown that putative quinone oxidoreductase complexes composed of proteins that contain a transmembrane diheme *b*-type cytochrome domain fused or adjacent to a multi-heme *c*-type cytochrome localized to the inner membrane are differentially expressed in response to changes in the reduction potential of the extracellular electron acceptor (1-4). These cytochrome *bc* (Cbc) complexes are highly conserved among *Geobacter* (5, 6) and are also present in the *G. metallireducens* genome (7).

Many of these Cbc complex genes were differentially expressed by *G. metallireducens* when grown in various co-culture conditions. The gene coding for CbcL (Gmet_0100) was ≥2.96 times more highly expressed in co-cultures with *G. sulfurreducens* than it was in any of the *Methanosarcina* co-cultures (Table 1). In *G. sulfurreducens*, CbcL was found to be important for electron transfer to extracellular electron acceptors with low reduction potentials such as insoluble Fe(III) oxide or an electrode poised at a potential of -0.1 V (3). Genes from another operon that codes for a Cbc quinone oxidoreductase complex composed of 5 proteins; 3 periplasmic *c*-type cytochromes *cbcD*, *cbcA* and *cbcC* (Gmet_2931, Gmet_2928, Gmet_2930), an inner membrane *b*-type cytochrome *cbcB* (Gmet_2929), and a membrane protein *cbcE* (Gmet_2932) were also much more highly expressed in co-cultures with *G.*

*sulfurreducens* than co-cultures with *Methanosarcina* (Table 1). Transcriptomic and genetic studies showed that the CbcABCDE complex is important for Fe(III) oxide reduction by *G. sulfurreducens* (8), however, deletion of *cbcA* and *cbcC* in *G. metallireducens* did not affect co-culture growth with any of the partners (this study).

CbcSTU is another Cbc complex that had high transcript abundance in *G. sulfurreducens* cells grown with Fe(III) oxide (8) that has two homologs in *G. metallireducens* (Gmet_0325-0327 and Gmet_3518-3520). Genes for both of the CbcSTU complexes were also more highly expressed by *G. metallireducens* cells grown in co-culture with *G. sulfurreducens* than in co-culture with *Methanosarcina* (Table 1), but they were not differentially expressed by *G. metallireducens* cells grown with Fe(III) oxide compared to Fe(III) citrate (9).

Another Cbc complex with a homolog in *G. sulfurreducens* consists of three proteins, a *b*-type cytochrome (*cbcW;* Gmet_1923), a 5-heme *c*-type cytochrome (*cbcX*; Gmet_1924) and an iron-sulfur cluster binding protein (*cbcV*; Gmet_1922) (8). Transcriptomic and genetic studies also showed that the CbcWXV complex was important for Fe(III) oxide reduction by *G. metallireducens*, *G. sulfurreducens*, and *G. uraniireducens* (8, 9), and genes from this complex were most highly expressed in co-cultures with *M. barkeri* (Table 2).

*G. metallireducens* cells grown with Type II *Methanosarcina* had high transcript abundance for genes from a Cbc complex consisting of a *b*-type cytochrome (*cbcP*; Gmet_0539), a transmembrane protein (*cbcQ*; Gmet_0533), an iron-sulfur cluster binding protein (*cbcO*; Gmet_0538), and three *c*-type cytochromes (*cbcM*; Gmet_0536), (*cbcN*; Gmet_0537) and (*cbcR*; Gmet_0534) (Table 3). Transcriptomic studies showed that genes from this complex were important for Fe(III) oxide reduction by *G. sulfurreducens* and *G. metallireducens* (8, 9), however, deletion of *cbcR* did not impact Fe(III) oxide reduction by *G. metallireducens* (9) and also did not affect co-culture growth with any of the partners (this study).

The *G. metallireducens* genome also has a cluster of genes (*pcmABCDEFG*) that code for another putative quinol-cytochrome *bc* complex without a homolog in *G. sulfurreducens* that is found in an operon with genes coding for proteins involved in para-cresol degradation (10-12). The PcmABCDEFG complex contains genes for an inner membrane protein (Gmet_2117), two inner membrane associated *b*-type cytochromes (*pcmC* and *pcmD*; Gmet_2119-2120), two iron sulfur cluster proteins (*pcmB*, *pcmE*; Gmet_2118, Gmet_2121), and two periplasmic *c*-type cytochrome proteins (*pcmF* and *pcmG*, Gmet_2122-2123) that were all more highly expressed in co-cultures with *M. barkeri* (Table 2).

**Other genes that were more highly expressed by *G. metallireducens* grown in co-culture with *G. sulfurreducens***

Transcript levels for genes for the flagellar filament protein, FliC, many of the flagellar biosynthesis proteins and the carbon storage regulator protein A (CsrA), which positively regulates flagellar synthesis (13), were most abundant in co-cultures with *G. sulfurreducens* (Supplementary Table S6). This high expression of *G. metallireducens* flagellar components during growth with *G. sulfurreducens* can be explained by the fact that *G. sulfurreducens* strain PCA does not produce a functional flagellum (14) and motility helps with the formation of aggregates. Flagella, however, have been observed on *M. acetivorans* (15). Although they are not directly involved in electron transfer, studies have shown that *Geobacter* flagella are important for growth on extracellular electron acceptors. Motility is needed for *Geobacter* to access insoluble electron acceptors, such as insoluble Fe(III) oxide, electrodes, or other organisms for participation in DIET. In fact, *G. metallireducens* strains lacking *fliC* were not capable of growth when insoluble Fe(III) oxide was provided as an electron acceptor (16) and it has been proposed that *Geobacter* flagella can serve as biofilm matrix scaffolds that enhance biofilm conductivity on current harvesting anodes (17). Deletion of *fliC* significantly increased the amount of time needed for establishment of aggregates with all DIET partners (this study). However, once aggregates became established, growth was similar to the wild-type co-cultures.

Consistent with the finding that many more genes coding for *c*-type cytochromes were more highly expressed by *G. metallireducens* grown in co-culture with *G. sulfurreducens* (Figure 2), genes coding for heme biogenesis proteins were also more highly expressed in the GM/GS co-culture (Supplementary Table S3). L-glutamate is a precursor for siroheme biosynthesis (18) and glutamate biosynthesis genes such as NADP dependent glutamate dehydrogenase (*gdhA*; Gmet_1186) and ferredoxin dependent glutamate synthase (*gltS*; Gmet_0147) were also much more highly expressed in GM/GS co-cultures than any of the *Methanosarcina* co-cultures (Supplementary Table S3C). In addition, genes coding for cysteine were more highly expressed in GM/GS co-cultures (Supplementary Table S3D), likely because synthesis of multi-heme cytochrome *c* holoproteins require high concentrations of cysteine because each heme group is covalently attached to two cysteine residues (19).

Genes involved in sulfate/sulfur assimilation and transport were among the most highly expressed genes in GM/GS co-cultures. In fact, genes for operons coding for sulfate/thiosulfate transport system proteins (*cysPTWA*; Gmet_1903-1906) and sulfate-sulfur assimilation proteins (*cysDN*; Gmet_2859-2860) were 42-731 and 4-15 times more highly expressed in GM/GS co-cultures than those with *Methanosarcina* (Supplementary Table S3E).

Transcript abundance for iron uptake and metabolism genes was also higher in GM/GS co-cultures. For example, the number of transcripts for the gene that codes for the iron transport protein, FeoB (Gmet_2444) was >3 times more highly expressed in GM/GS co-cultures than those with *Methanosarcina* (Supplementary Table S3F).

Genes involved in synthesis of ferredoxin were more highly expressed in GM/GS co-cultures. The genes for cysteine desulfurase (*iscS*; Gmet_0872) and its transcriptional regulator (*iscR*; Gmet_0871) were 10 to 36 times more highly transcribed in GM/GS co-cultures (Supplementary Table S3G). In addition, the gene coding for the chaperone involved in maturation of iron-sulfur cluster containing proteins, HscA (Gmet_3532) and numerous genes coding for proteins with iron-sulfur binding domains were more highly expressed in GM/GS co-cultures (Supplementary Table S2).

Transcriptomic analyses revealed that the composition of the *G. metallireducens* cell membrane was slightly different when it was grown via DIET with *G. sulfurreducens* compared to growth with any of the *Methanosarcina* species. *G. metallireducens* membranes are known to contain hopanoid lipids (20), which are important for bacterial membrane stability and functioning (21). Genes for hopene biosynthesis were more highly expressed in GM/GS co-cultures (Supplementary Table S3H). Studies have shown that the presence of the hopanoid lipid, hopene, lowers membrane ion permeability, membrane fluidity and membrane conductance (20, 22) and this might be important for electron transfer to other *Geobacter* species. However, further investigation into this possibility is required.

**Differences between Type I and Type II *Methanosarcina***

Aside from differences in quinol oxidoreductase complex proteins, many of the electron transport genes more highly expressed in GM/MB co-cultures than in other co-culture conditions coded for periplasmic *c*-type cytochromes (Table 2). These included Gmet_1703 and Gmet_2156, which code for periplasmic 7- and 9-heme cytochromes, a periplasmic tri-heme cytochrome, *ppcF* (Gmet_0335), and 5 periplasmic di-heme cytochromes, *ppcB* (Gmet_3166), *ccpB* (Gmet_1210), *ppcE* (Gmet_1846), *narC* (Gmet_0328), and *ppcA* (Gmet_2902). The only one of these *c*-type cytochromes that was more highly expressed by *G. metallireducens* during growth on Fe(III) oxide was Gmet_2156 (9). A *ccpB* homolog in *G. sulfurreducens* codes for a cytochrome *c* peroxidase protein that protects the cell from reactive oxygen species (23), and Ppc cytochromes are periplasmic electron transporters involved in Fe(III) respiration by *G. sulfurreducens* (24, 25).

Genes coding for several *c*-type cytochrome proteins were also more highly expressed in Type II *Methanosarcina* compared to *M. barkeri* (Table 3). These included a periplasmic monoheme cytochrome (Gmet_0232), the periplasmic di-heme cytochrome peroxidase CcpA (Gmet_3091) (23), and Gmet_0679, which codes for a 5-heme cytochrome with unknown localization. Gmet_0679 was highly expressed in Fe(III) oxide grown *G. metallireducens* cells, but gene deletion studies revealed that it was not essential for Fe(III) oxide reduction (9). Deletion of Gmet_0679 and Gmet_0232 from *G. metallireducens* also did not impact growth via DIET with any of the electron-accepting partners (this study).

Genes coding for proteins involved in the biosynthesis of menaquinone and ubiquinone (26, 27), lipid soluble electron carriers involved in electron shuttling across the inner membrane, were most highly expressed by *G. metallireducens* grown via DIET with Type II *Methanosarcina* (Supplementary Table S3I). The gene coding for UbiA, which codes for a 4-hydroxy benzoate octaprenyl transferase thought to be a key enzyme of ubiquinone biosynthesis (26) was more than ~4 times more highly expressed in Type II *Methanosarcina* co-cultures.

Genes for biosynthesis of the coenzymes cobalamin, folate, molybdopterin, and thiamine were more significantly expressed by *G. metallireducens* in Type II *Methanosarcina* co-cultures (Supplementary Table S3J-M). *Methanosarcina* make several methanogenesis pathway enzymes that require such cofactors (28-30) and studies have shown that cross-feeding of vitamins like cobalamin are common in interspecies symbiotic partnerships (31-34). It is possible that *G. metallireducens* makes higher concentrations of these coenzymes in the presence of Type II *Methanosarcina* because it participates in higher rates of cross-feeding with these methanogens than with the other electron accepting partners. Evidence for increased cross-feeding to Type II *Methanosarcina* is provided by the finding that expression of *btuB* (Gmet_1245) which codes for an outer membrane cobalamin transporter (35), was >5 and >23 fold higher in Type II *Methanosarcina*-GM co-cultures than GM/GS and GM/MB co-cultures (Supplementary Table S2).

**Similarities in expression of non-cytochrome genes during DIET**

A gene just upstream from Gmet_0930, Gmet_0933, was among the most highly expressed genes in all 4 conditions (Supplementary Table S1). Gmet_0933 codes for a protein that shares many similarities to Bap (biofilm-associated protein) proteins, which are large secreted proteins required for biofilm formation in various bacteria (36, 37). Similar to previously characterized Bap proteins, Gmet_0933 has a number of tandem repeats (24 repeats), contains two cadherin like domains, an EF-hand calcium binding motif, a serine rich region, and 6 immunoglobulin like domains. Gmet_0933 also contains 7 mirror beta grasp (MBG) domains that are associated with bacterial surface proteins (38). Another gene (Gmet_2043) that codes for a large and highly repetitive (15 tandem repeats) outer surface associated protein that is potentially involved in biofilm formation was also highly transcribed by *G metallireducens* in all 4 co-culture conditions (Supplementary Table S1). It has five SdrD (SD-repeat containing protein D) B-like domains, which are found in cell surface adhesion proteins (39).

The gene coding for OmpJ (Gmet_3254) was the most highly expressed membrane associated protein in all 4 co-cultures (Supplementary Table S1). It is homologous to an outer membrane porin found in *G. sulfurreducens* that is indirectly required for Fe(III) oxide reduction (40). It serves as a putative porin that helps stabilize the integrity of the periplasmic space necessary for proper folding and functioning of periplasmic and outer membrane electron transport components. Another gene coding for an outer membrane porin protein, OmpA (Gmet_0342) was also among the most highly expressed genes in all 4 conditions. The gene coding for OmpA was also highly expressed by *G. sulfurreducens* and *G. metallireducens* cells during growth in the presence of Fe(III) (9, 41), and it has been shown to play a role in biofilm formation (42).

Almost all of the genes from an operon (Gmet_2022-2033) that codes for proteins involved in production of exopolysaccharides and lipopolysaccharides were also highly expressed by *G. metallireducens* in all 4 conditions. Lipopolysaccharides (LPS) are important for bacterial adherence to surfaces during biofilm formation in many Gram-negative bacteria (43). Transcriptomic and genetic studies showed that Gmet_2029 which codes for a putative lipopolysaccharide synthesis protein was critical for Fe(III) oxide reduction (9), and it was important for DIET in all 4 co-cultures. In fact, *G. metallireducens* strains in which Gmet_2029 was deleted were unable to form aggregates with *G. sulfurreducens*. Establishment of co-culture growth with *Methanosarcina* species was significantly delayed and once co-cultures were established they continued to grow 2.2 ($p=0.002$), 2.81 ($p=0.0003$), and 3.4 ($p=0.009$) times slower in ΔGmet_2029/MB, ΔGmet_2029/MA, and ΔGmet_2029/MS co-cultures than the wild-type co-cultures after 4 transfers (this study).

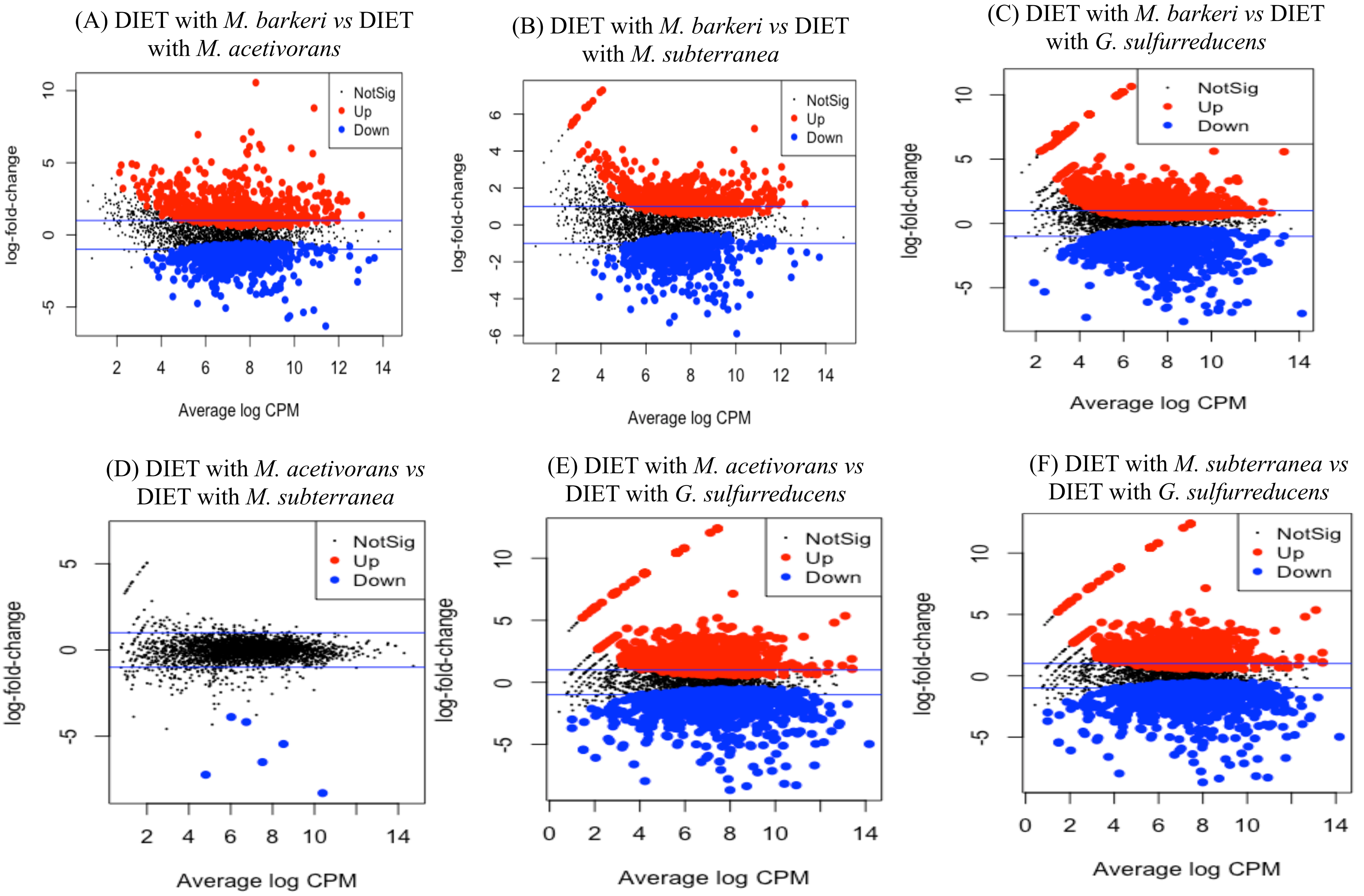


Supplementary Figure S1. Mean-difference plots (MA plots) generated with plotMD from the LIMMA package in RStudio comparing *G. metallireducens* normalized transcripts from various co-cultures. Genes that were significantly up-regulated are highlighted in red while those that were significantly down-regulated are highlighted in blue. All plots were generated from triplicate RNAseq libraries. Plots A, B, and C show differentially expressed genes from *M. barkeri* co-cultures; Plots D and E show differentially expressed genes from *M. acetivorans* co-cultures; Plot F shows differentially expressed genes from *M. subterranea* co-culture.

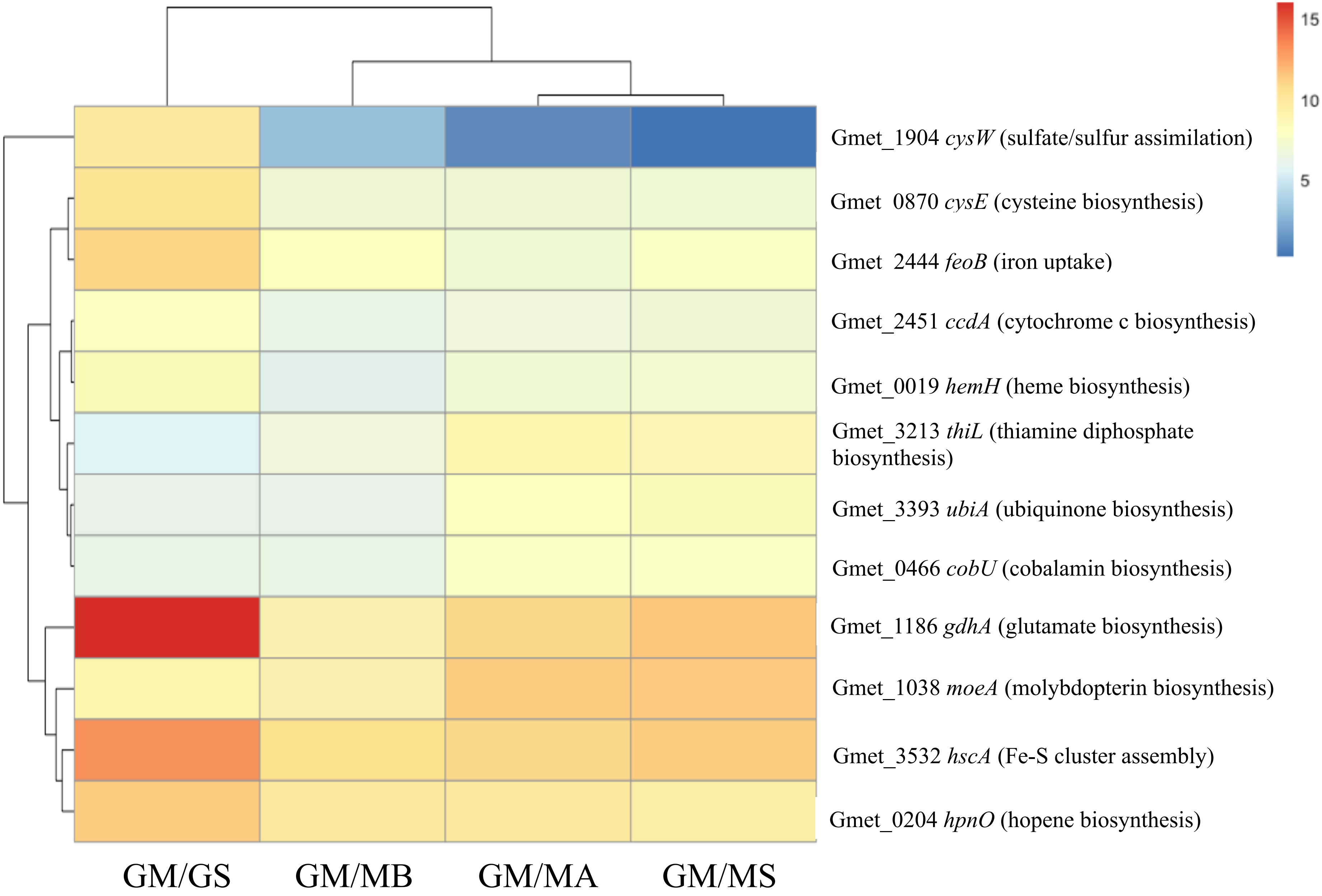


Supplementary Figure S2. $Log_2$ read values normalized with the TMM method for representative biosynthetic pathway genes in *G. metallireducens* from 4 different RNAseq co-culture libraries (GM/GS: *G. metallireducens* and *G. sulfurreducens*, GM/MB: *G. metallireducens* and *M. barkeri*, GM/MA: *G. metallireducens* and *M. acetivorans*, GM/MS: *G. metallireducens* and *M. subterranea*) that had $log_2$ values above the median.

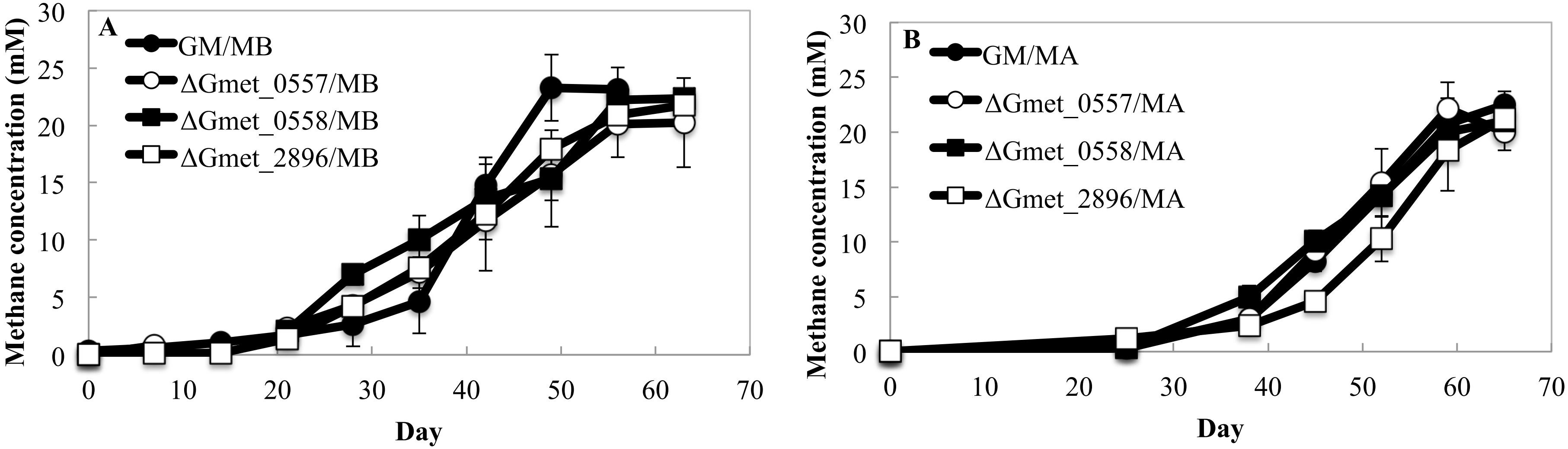


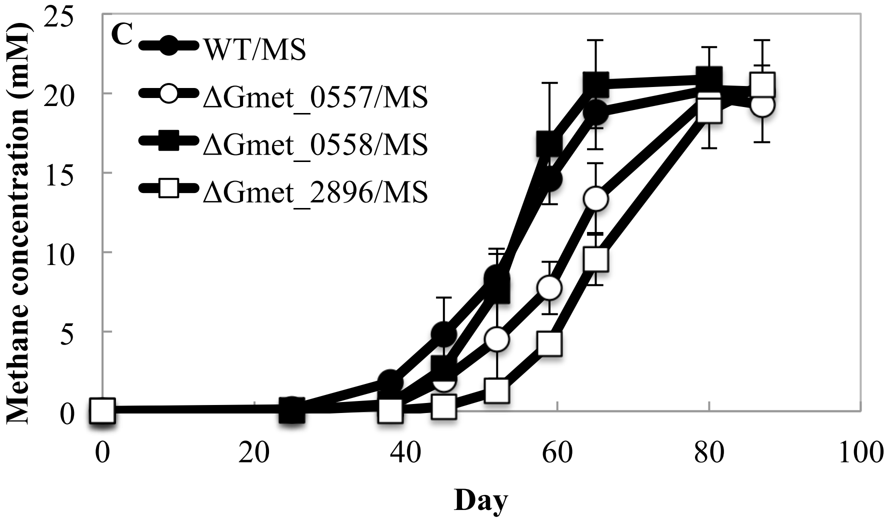


Supplementary Figure S3. Methane production by various co-cultures during the first transfer when aggregates first became established. (A) *G. metallireducens* strains and *M. barkeri* (B) *G. metallireducens* strains and *M. acetivorans*; (C) *G. metallireducens* strains and *M. subterranea*. Ethanol (20 mM) was provided as the electron donor for all co-cultures and error bars represent the standard deviation between triplicate cultures.

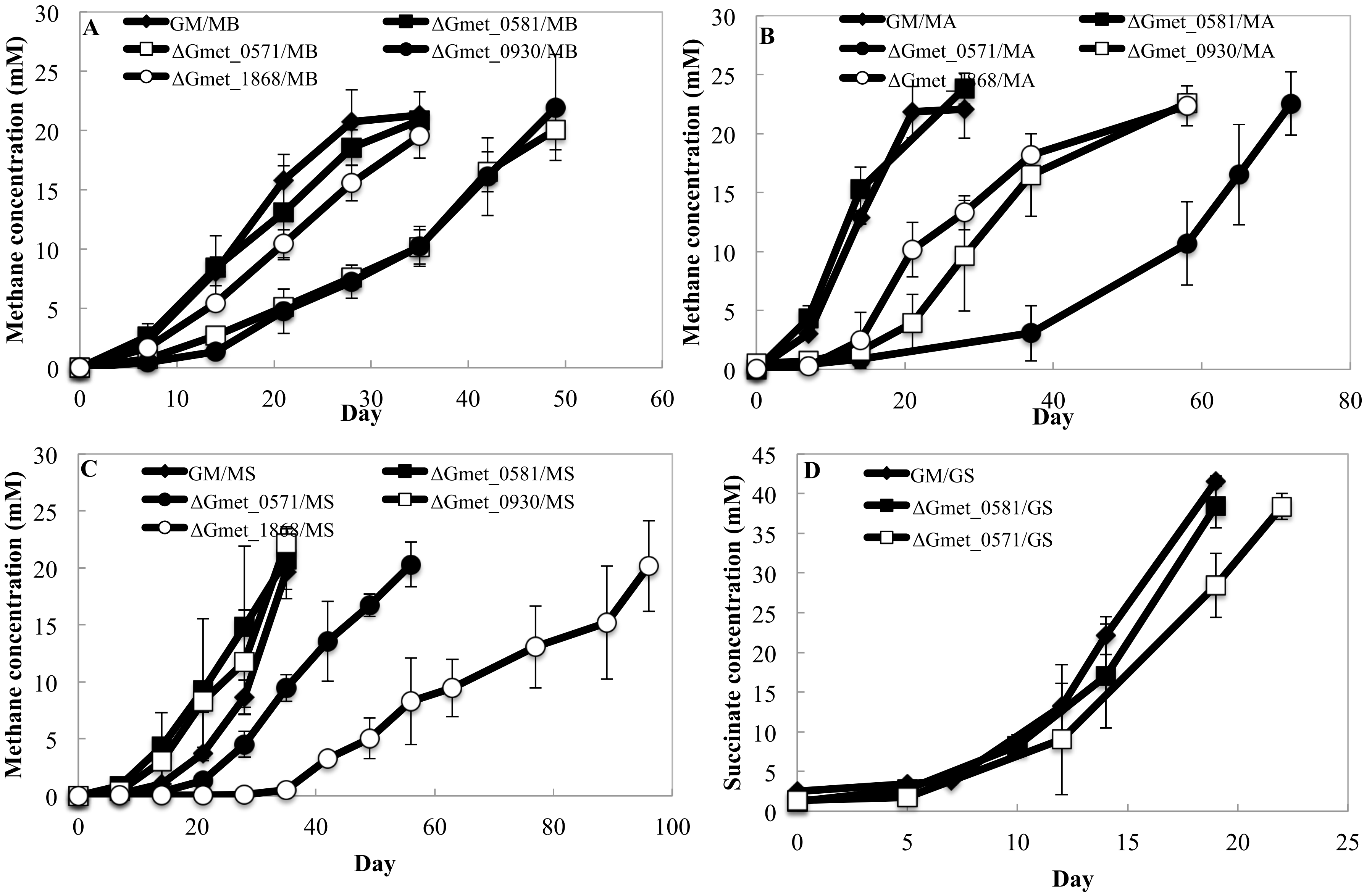


Supplementary Figure S4. Methane or succinate production by co-cultures with various species of *Methanosarcina* or *G. sulfurreducens* and *G. metallireducens* strains lacking the c-type cytochromes Gmet_0571, Gmet_0581, Gmet_1868, and Gmet_0930 after 4 transfers. (A) Co-cultures with *M. barkeri*; (B) Co-cultures with *M. acetivorans*; (C) Co-cultures with *M. subterranea*. (D) Co-cultures with *G. sulfurreducens*. Error bars represent the standard deviation from triplicate cultures. Growth was 1.7 (p= 0.005), 2.0 (p=0.001), and 1.6 (p=0.009) times slower in ΔGmet_0930/MB, ΔGmet_0930/MA, and ΔGmet_0930/MS co-cultures than wild-type; growth was 1.96 (p=0.003), 3.04 ($p=7.56x10^{-5}$), and 1.6 (p=0.04) times slower in ΔGmet_0571/MB, ΔGmet_0571/MA and ΔGmet_0571/MS co-cultures than wild-type; growth was 1.4 (p=0.03), 1.6 (p=0.001), and 3.3 (p=0.001) times slower in GM/MB, GM/MA, and GM/MS co-cultures than wild-type.

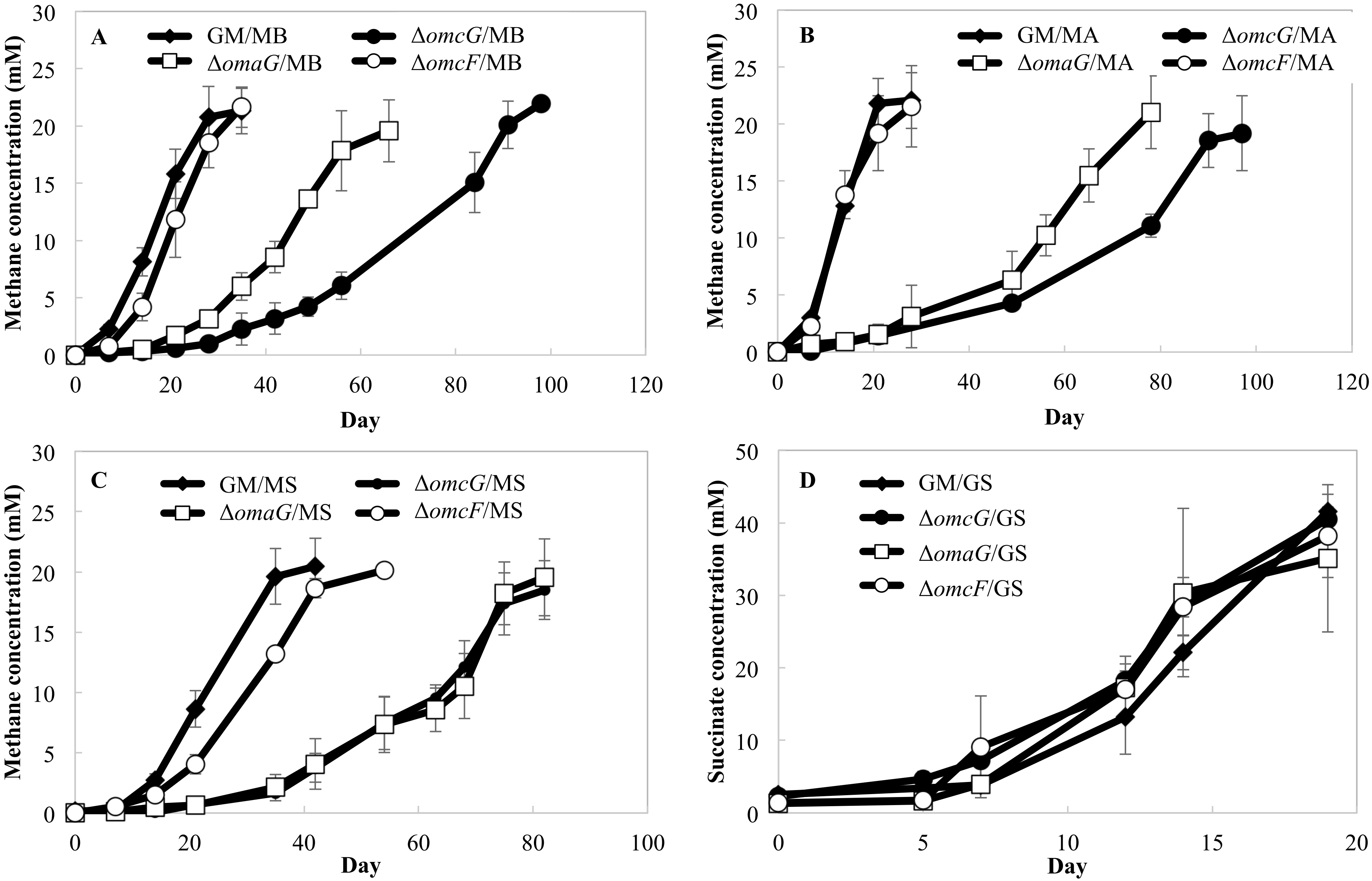


Supplementary Figure S5. Methane or succinate production by co-cultures with various *Methanosarcina* species or *G. sulfurreducens* and either wild-type *G. metallireducens* or the *omcG, omcF*, or the *omaG* deletion mutant strain during the 4[th] transfer. (A) Co-cultures with *M. barkeri*; (B) Co-cultures with *M. acetivorans*; (C) Co-cultures with *M. subterranea*. (D) Co-cultures with *G. sulfurreducens.* Error bars represent the standard deviation from triplicate cultures. Even after 4 transfers, the Δ*omcG*/MB, Δ*omcG*/MA, and Δ*omcG*/MS co-cultures grew 3.64 ($p=2.62\times10^{-5}$), 4.58 ($p=0.0005$), and 2.3 ($p=0.003$) times slower than wild-type-*Methanosarcina* co-cultures, and Δ*omaG*/MB, Δ*omaG*/MA, and Δ*omaG*/MS co-cultures grew 2.21 ($p=0.001$), 3.59 ($p=0.0007$), and 3.79 ($p=0.0006$) times slower than wild-type-*Methanosarcina* co-cultures.

Supplementary Table S3A. Average normalized TMM $\log_2$ values for *G. metallireducens* genes involved in the biosynthesis of *c*-type cytochromes

| Locus ID | Gene | GM/MB | GM/MA | GM/MS | GM/GS |
|---|---|---|---|---|---|
| **Gmet_0573** | *ccsB* | 5.95 | 6.89 | 5.31 | 7.65 |
| **Gmet_3232** | *ccsC* | 7.64 | 7.99 | 8.12 | 9.15 |
| **Gmet_2900** | *ccsAB* | 8.32 | 8.58 | 8.69 | 10.80 |
| **Gmet_2451** | *ccdA-1* | 6.24 | 6.83 | 6.91 | 7.86 |
| **Gmet_1384** | *ccdA-2* | 4.93 | 2.83 | 3.03 | 5.14 |
| **Gmet_2452** | *resA* | 7.15 | 6.66 | 7.00 | 6.94 |
| **Gmet_2901** | *resB* | 10.21 | 9.42 | 9.49 | 11.04 |
| **Gmet_0579** | *ccsB* | 5.32 | 5.38 | 5.28 | 5.61 |
| **Gmet_0594** | *ccsB* | 6.65 | 6.62 | 6.71 | 7.11 |
| **Gmet_0141** | *ccsA* | 5.93 | 6.43 | 6.32 | 6.18 |
| | Average | 6.83 | 6.76 | 6.68 | 7.75 |

Supplementary Table S3B. Average normalized TMM $\log_2$ values for *G. metallireducens* genes involved in heme biosynthesis

| Locus ID | Gene | GM/MB | GM/MA | GM/MS | GM/GS |
|---|---|---|---|---|---|
| **Gmet_0015** | | 9.28 | 9.61 | 10.08 | 11.84 |
| **Gmet_0016** | *hemE* | 8.89 | 9.81 | 9.81 | 11.65 |
| **Gmet_0189** | *hemB* | 8.87 | 9.18 | 9.62 | 10.89 |
| **Gmet_3234** | *hemC* | 8.43 | 9.43 | 9.21 | 10.50 |
| **Gmet_3233** | *hemA* | 8.66 | 9.26 | 9.23 | 9.94 |
| **Gmet_3551** | *hemY* | 9.42 | 9.75 | 10.16 | 9.94 |
| **Gmet_3356** | *hemL* | 9.49 | 8.67 | 9.03 | 9.79 |
| **Gmet_3235** | *hemD* | 8.31 | 9.73 | 9.78 | 9.18 |
| **Gmet_1736** | *gltX* | 7.88 | 8.03 | 8.17 | 9.00 |
| **Gmet_2750** | *hemE* | 5.55 | 8.30 | 8.07 | 8.98 |
| **Gmet_0476** | *cbiX* | 6.34 | 6.86 | 6.51 | 8.86 |
| **Gmet_0019** | *hemH* | 5.94 | 7.14 | 7.18 | 8.41 |
| **Gmet_3404** | *pdxA* | 5.58 | 6.68 | 7.10 | 7.62 |
| **Gmet_0253** | *ctaB* | 7.38 | 7.25 | 6.68 | 6.72 |
| **Gmet_3535** | *hemN* | 6.89 | 7.79 | 8.16 | 6.36 |
| **Gmet_3231** | *cysG* | 6.10 | 7.16 | 7.15 | 5.87 |
| | Average | 7.69 | 8.41 | 8.50 | 9.10 |

Supplementary Table S3C. Average normalized TMM $\log_2$ values for *G. metallireducens* genes involved in L-glutamate biosynthesis

| Locus ID | Gene | GM/MB | GM/MA | GM/MS | GM/GS |
|---|---|---|---|---|---|
| **Gmet_1186** | *gdhA* | 8.90 | 10.92 | 11.58 | 16.01 |
| **Gmet_0147** | *gltS* | 7.99 | 9.35 | 8.90 | 10.45 |
| **Gmet_1940** | | ND | ND | ND | ND |
| **Gmet_1728** | | 8.31 | 8.73 | 8.71 | 9.00 |
| | Average | 8.40 | 9.67 | 9.73 | 11.82 |

Supplementary Table S3D. Average normalized TMM $log_2$ values for *G. metallireducens* genes involved in L-cysteine and L-methionine biosynthesis

| Locus ID | Gene | GM/MB | GM/MA | GM/MS | GM/GS |
|---|---|---|---|---|---|
| **Gmet_0870** | *cysE* | 6.90 | 7.00 | 7.09 | 10.30 |
| **Gmet_1558** | *cysK-2* | 8.44 | 9.26 | 9.53 | 10.21 |
| **Gmet_2988** | *cysK-1* | 8.47 | 10.03 | 9.94 | 13.06 |
| **Gmet_0264** | *cysM* | 7.03 | 8.51 | 8.48 | 8.14 |
| **Gmet_0699** | *metC-2* | 6.21 | 7.22 | 6.88 | 11.86 |
| | Average | 7.41 | 8.40 | 8.38 | 10.71 |

Supplementary Table S3E. Average normalized TMM $log_2$ values for *G. metallireducens* genes involved in sulfur metabolism

| Locus ID | Gene | GM/MB | GM/MA | GM/MS | GM/GS |
|---|---|---|---|---|---|
| **Gmet_2520** | | 6.67 | 13.04 | 6.22 | 11.20 |
| **Gmet_1904** | *cysW* | 3.06 | 1.04 | 0.20 | 9.93 |
| **Gmet_1906** | *cysP* | 4.08 | 4.26 | 4.54 | 9.92 |
| **Gmet_1903** | *cysA* | 2.80 | 1.92 | 3.12 | 9.85 |
| **Gmet_1905** | *cysU* | 3.35 | 2.97 | 2.64 | 9.47 |
| **Gmet_2857** | | 4.80 | 6.35 | 4.90 | 9.07 |
| **Gmet_1588** | | 8.56 | 8.06 | 8.08 | 9.06 |
| **Gmet_2860** | *cysN* | 5.81 | 6.81 | 5.67 | 8.99 |
| **Gmet_2859** | *cysD* | 5.55 | 6.22 | 5.03 | 8.87 |
| **Gmet_0531** | | 7.64 | 7.68 | 7.49 | 8.47 |
| **Gmet_2858** | *cysH* | 4.75 | 5.55 | 4.01 | 8.34 |
| **Gmet_3431** | *hdrF* | 6.30 | 7.50 | 8.22 | 8.17 |
| **Gmet_3430** | *hdrE* | 6.67 | 8.02 | 8.34 | 7.58 |
| **Gmet_0998** | | 6.00 | 5.49 | 5.31 | 5.26 |
| **Gmet_0996** | | 6.66 | 5.69 | 5.06 | 3.39 |
| **Gmet_0997** | | 5.92 | 5.34 | 4.95 | 3.35 |
| | Average | 5.54 | 6.00 | 5.24 | 8.18 |

Supplementary Table S3F. Average normalized TMM $log_2$ values for *G. metallireducens* genes involved in iron uptake and metabolism

| Locus ID | Gene | GM/MB | GM/MA | GM/MS | GM/GS |
|---|---|---|---|---|---|
| **Gmet_2444** | *feoB* | 8.05 | 7.12 | 7.73 | 11.04 |
| **Gmet_2442** | *feoA* | 6.11 | 6.26 | 6.16 | 9.04 |
| **Gmet_2445** | *fur* | 10.32 | 9.89 | 10.79 | 9.02 |
| | Average | 8.16 | 7.75 | 8.23 | 9.70 |

Supplementary Table S3G. Average normalized TMM $log_2$ values for *G. metallireducens* genes involved in iron-sulfur cluster biosynthesis

| Locus ID | Gene | GM/MB | GM/MA | GM/MS | GM/GS |
|---|---|---|---|---|---|
| **Gmet_0871** | *iscR* | 5.16 | 4.27 | 4.40 | 9.56 |
| **Gmet_0872** | *iscS* | 6.20 | 6.35 | 6.35 | 9.79 |
| **Gmet_0991** | *iscU* | 11.86 | 11.94 | 11.79 | 12.39 |
| **Gmet_0992** | *iscS* | 11.65 | 12.52 | 12.46 | 11.95 |
| **Gmet_1300** | *iscS* | 8.68 | 9.50 | 9.47 | 8.51 |
| **Gmet_1907** | *iscR* | 6.91 | 6.45 | 6.74 | 9.24 |
| **Gmet_2006** | *sudS* | 8.66 | 9.13 | 9.61 | 7.23 |
| **Gmet_2934** | *thiG* | 6.38 | 7.93 | 8.07 | 8.40 |
| **Gmet_3532** | *hscA* | 10.63 | 10.85 | 11.23 | 13.02 |
| | Average | 8.46 | 8.77 | 8.90 | 10.01 |

Supplementary Table S3H. Average normalized TMM $log_2$ values for G. *metallireducens* genes involved in hopene biosynthesis

| Locus ID | Gene | GM/MB | GM/MA | GM/MS | GM/GS |
|---|---|---|---|---|---|
| **Gmet_2971** | *hpnP* | 8.44 | 9.55 | 10.09 | 10.47 |
| **Gmet_0204** | *hpnO* | 9.85 | 9.82 | 9.30 | 11.14 |
| **Gmet_1274** | *hpnG* | 6.70 | 6.95 | 6.95 | 6.06 |
| **Gmet_2823** | *hpnH* | 9.30 | 8.47 | 9.22 | 11.32 |
| **Gmet_0419** | *hpnF* | 6.56 | 6.65 | 6.53 | 8.01 |
| **Gmet_2820** | *hpnF* | 10.12 | 10.34 | 10.29 | 10.79 |
| **Gmet_2821** | *hpnA* | 7.54 | 8.75 | 8.45 | 8.09 |
| | Average | 8.36 | 8.65 | 8.69 | 9.41 |

Supplementary Table S3I. Average normalized TMM $log_2$ values for *G. metallireducens* genes involved in ubiquinone and menaquinone biosynthesis

| Locus ID | Gene | GM/MB | GM/MA | GM/MS | GM/GS |
|---|---|---|---|---|---|
| **Gmet_3392** | *ubiX* | 5.57 | 6.78 | 7.10 | 5.73 |
| **Gmet_3380** | *mqnD* | 6.06 | 7.43 | 6.87 | 6.71 |
| **Gmet_3393** | *ubiA* | 6.18 | 8.14 | 8.34 | 6.19 |
| **Gmet_3391** | *mqnE* | 6.86 | 7.86 | 8.01 | 7.97 |

| | | | | | |
|---|---|---|---|---|---|
| **Gmet_3381** | *mqnB* | 7.31 | 8.41 | 7.80 | 7.15 |
| **Gmet_3390** | *mqnC-1* | 7.91 | 8.59 | 8.72 | 8.98 |
| **Gmet_0986** | *mqnA* | ND | ND | ND | ND |
| **Gmet_0993** | *ubiD* | ND | ND | ND | ND |
| **Gmet_1167** | *ubiE* | ND | ND | ND | ND |
| **Gmet_1645** | | ND | ND | ND | ND |
| **Gmet_0771** | *wrbA* | ND | ND | ND | ND |
| | Average | 6.65 | 7.87 | 7.80 | 7.12 |

Supplementary Table S3J. Average normalized TMM $\log_2$ values for *G. metallireducens* genes involved in cobalamin biosynthesis

| Locus ID | Gene | GM/MB | GM/MA | GM/MS | GM/GS |
|---|---|---|---|---|---|
| **Gmet_0466** | *cobU* | 6.23 | 7.64 | 7.65 | 6.26 |
| **Gmet_0467** | *cobT* | 7.59 | 8.85 | 8.36 | 9.08 |
| **Gmet_0468** | *cobS* | 5.96 | 8.45 | 7.88 | 7.29 |
| **Gmet_0469** | *cobC* | 5.68 | 7.20 | 6.30 | 6.00 |
| **Gmet_0477** | *cbiC* | 5.77 | 8.05 | 7.38 | 5.80 |
| **Gmet_0478** | *cbiD* | 8.10 | 9.79 | 9.48 | 7.32 |
| **Gmet_0481** | *cbiET* | 7.79 | 8.99 | 8.60 | 10.49 |
| **Gmet_0484** | *cbiG* | 7.36 | 8.75 | 8.84 | 6.81 |
| **Gmet_0485** | *cbiH_cbiP* | 7.69 | 9.04 | 9.06 | 9.05 |
| **Gmet_0486** | *cbiB* | 5.56 | 7.09 | 6.91 | 4.21 |
| **Gmet_1573** | *cobA* | 8.80 | 6.42 | 6.56 | 7.76 |
| | Average | 6.96 | 8.21 | 7.91 | 7.28 |

Supplementary Table S3K. Average normalized TMM $\log_2$ values for *G. metallireducens* genes involved in folate biosynthesis

| Locus ID | Gene | GM/MB | GM/MA | GM/MS | GM/GS |
|---|---|---|---|---|---|
| **Gmet_1162** | *folD-2* | 8.50 | 8.72 | 8.93 | 10.14 |
| **Gmet_1605** | *glyA* | 8.80 | 9.62 | 9.77 | 8.98 |
| **Gmet_1657** | *queD* | 7.80 | 6.95 | 6.65 | 7.73 |
| **Gmet_1889** | *folP* | 8.30 | 9.42 | 8.80 | 7.03 |
| **Gmet_2475** | *folC* | 8.13 | 8.94 | 8.80 | 7.37 |
| **Gmet_2947** | *folA* | 3.35 | 6.48 | 6.46 | 4.68 |
| **Gmet_3010** | | 7.74 | 9.27 | 9.52 | 7.47 |
| **Gmet_3074** | *folE* | 6.88 | 7.37 | 6.93 | 8.21 |
| **Gmet_3123** | | 5.66 | 8.02 | 8.01 | 5.06 |

| | | | | | |
|---|---|---|---|---|---|
| **Gmet_0247** | *folD-1* | 8.77 | 10.07 | 9.89 | 9.92 |
| | Average | 7.39 | 8.49 | 8.38 | 7.66 |

Supplementary Table S3L. Average normalized TMM $\log_2$ values for *G. metallireducens* genes involved in molybdopterin biosynthesis

| Locus ID | Gene | GM/MB | GM/MA | GM/MS | GM/GS |
|---|---|---|---|---|---|
| **Gmet_0301** | *moaA-1* | 7.31 | 8.08 | 8.37 | 9.70 |
| **Gmet_0336** | *moeA-2* | 6.47 | 7.33 | 7.33 | 4.46 |
| **Gmet_0337** | *moaA-2* | 5.74 | 6.39 | 6.03 | 9.05 |
| **Gmet_1036** | *moaB* | 6.69 | 10.12 | 10.23 | 6.83 |
| **Gmet_1037** | *moaC* | 8.40 | 10.74 | 11.60 | 7.94 |
| **Gmet_1038** | *moeA-1* | 8.91 | 11.14 | 11.31 | 8.90 |
| **Gmet_1804** | *moeA-3* | 6.45 | 6.33 | 6.34 | 4.48 |
| **Gmet_2095** | *moaA-3* | 6.29 | 5.63 | 6.06 | 4.86 |
| | Average | 7.03 | 8.22 | 8.41 | 7.03 |

Supplementary Table S3M. Average normalized TMM $\log_2$ values for *G. metallireducens* genes involved in thiamine biosynthesis

| Locus ID | Gene | GM/MB | GM/MA | GM/MS | GM/GS |
|---|---|---|---|---|---|
| **Gmet_2909** | *thiE_thiD* | 8.61 | 9.51 | 9.32 | 9.05 |
| **Gmet_2910** | *thiC-1* | 8.14 | 8.92 | 8.89 | 10.56 |
| **Gmet_2935** | *tenI* | 6.15 | 7.45 | 7.58 | 6.15 |
| **Gmet_3213** | *thiL* | 6.81 | 8.87 | 8.59 | 5.70 |
| | Average | 7.43 | 8.69 | 8.59 | 7.86 |

Supplementary Table S3N. Average normalized TMM $\log_2$ values for *G. metallireducens* genes involved in riboflavin biosynthesis

| Locus ID | Gene | GM/MB | GM/MA | GM/MS | GM/GS |
|---|---|---|---|---|---|
| **Gmet_1388** | *ribF* | 6.85 | 7.14 | 6.95 | 6.26 |
| **Gmet_1624** | *ribD* | 7.17 | 7.47 | 6.98 | 7.44 |
| **Gmet_1625** | *ribE* | 5.89 | 6.38 | 6.04 | 5.71 |
| **Gmet_1626** | *ribA* | 10.20 | 9.59 | 9.52 | 10.96 |
| **Gmet_1627** | *ribH* | 7.06 | 8.33 | 7.81 | 6.40 |
| | average | 7.43 | 7.78 | 7.46 | 7.36 |

Supplementary Table S3O. Average normalized TMM $\log_2$ values for *G. metallireducens* genes involved in lipopolysaccharide biosynthesis

| Locus ID | Gene | GM/MB | GM/MA | GM/MS | GM/GS |
|---|---|---|---|---|---|

| | | | | | |
|---|---|---|---|---|---|
| **Gmet_1704** | *pagL* | 5.20 | 5.09 | 5.08 | 2.80 |
| **Gmet_0585** | *waaQ* | 5.62 | 6.19 | 6.53 | 4.53 |
| **Gmet_0920** | *gmhA* | 6.11 | 7.81 | 7.39 | 7.56 |
| **Gmet_0923** | *gmhB* | 6.16 | 7.03 | 7.39 | 5.67 |
| **Gmet_2331** | *waaC* | 6.31 | 7.14 | 7.38 | 8.75 |
| **Gmet_2347** | *lpxK* | 6.53 | 6.98 | 7.00 | 5.92 |
| **Gmet_2350** | *lpxB* | 6.74 | 7.39 | 7.14 | 8.11 |
| **Gmet_2345** | *waaF* | 6.86 | 7.47 | 7.72 | 5.81 |
| **Gmet_2341** | *waaQ* | 6.95 | 7.01 | 7.12 | 6.34 |
| **Gmet_2348** | *waaA* | 6.95 | 7.53 | 7.73 | 7.04 |
| **Gmet_1279** | *kdsC* | 7.52 | 6.33 | 6.82 | 7.10 |
| **Gmet_2776** | *lpxH* | 7.74 | 7.59 | 7.76 | 7.58 |
| **Gmet_1278** | *kdsD* | 7.81 | 8.57 | 8.62 | 6.99 |
| **Gmet_2607** | *lpxC* | 7.99 | 9.54 | 9.17 | 8.61 |
| **Gmet_1275** | *kdsB* | 8.10 | 8.26 | 8.40 | 7.30 |
| **Gmet_0091** | | 8.20 | 9.08 | 8.90 | 8.46 |
| **Gmet_2353** | *lpxA-1* | 8.44 | 9.06 | 9.11 | 8.78 |
| **Gmet_0922** | *hldE* | 8.45 | 8.76 | 8.93 | 8.19 |
| **Gmet_2567** | *lpxA-2* | 8.74 | 8.21 | 7.97 | 8.60 |
| **Gmet_1254** | *uppS* | 8.74 | 8.05 | 8.32 | 8.42 |
| **Gmet_1277** | *kdsA* | 8.89 | 9.14 | 9.21 | 7.90 |
| **Gmet_2355** | *lpxD* | 9.14 | 10.03 | 9.85 | 9.28 |
| **Gmet_2348** | *kdtA* | ND | ND | ND | ND |
| | Average | 7.42 | 7.83 | 7.89 | 7.26 |

Supplementary Table S3P. Average normalized TMM $\log_2$ values for *G. metallireducens* genes involved in peptidoglycan biosynthesis

| **Locus ID** | **Gene** | **GM/MB** | **GM/MA** | **GM/MS** | **GM/GS** |
|---|---|---|---|---|---|
| **Gmet_0690** | *uppP* | 4.50 | 2.06 | 1.17 | 1.80 |
| **Gmet_3403** | | 7.02 | 6.28 | 6.22 | 8.87 |
| **Gmet_2778** | | 7.72 | 8.26 | 7.92 | 7.92 |
| **Gmet_0382** | *murA* | 8.41 | 9.89 | 9.68 | 10.29 |
| **Gmet_0928** | *mrdA* | 8.61 | 7.82 | 8.21 | 7.54 |
| **Gmet_0409** | *mraY* | 8.68 | 9.52 | 9.78 | 8.90 |
| **Gmet_3133** | *uppP* | 8.75 | 7.98 | 7.53 | 7.37 |
| **Gmet_1671** | | 8.98 | 9.07 | 8.90 | 8.49 |
| **Gmet_0354** | *mrcA* | 9.27 | 9.45 | 9.72 | 10.41 |
| **Gmet_0408** | *murF* | 9.85 | 10.28 | 10.58 | 9.09 |
| **Gmet_0412** | *murG* | 9.95 | 9.80 | 10.23 | 9.63 |
| **Gmet_0407** | *murE* | 10.25 | 10.37 | 10.78 | 10.82 |

| | | | | | |
|---|---|---|---|---|---|
| **Gmet_0406** | *ftsI* | 10.92 | 10.86 | 11.12 | 10.44 |
| **Gmet_0410** | *murD* | ND | ND | ND | ND |
| **Gmet_0413** | *murC* | ND | ND | ND | ND |
| **Gmet_0414** | *murB* | ND | ND | ND | ND |
| **Gmet_0415** | *ddl_ftsQ* | ND | ND | ND | ND |
| | Average | 8.69 | 8.59 | 8.60 | 8.58 |

Supplementary Table S3Q. Average normalized TMM $\log_2$ values for *G. metallireducens* genes involved in fatty acid biosynthesis

| Locus ID | Gene | GM/MB | GM/MA | GM/MS | GM/GS |
|---|---|---|---|---|---|
| **Gmet_1469** | | 6.29 | 6.37 | 5.90 | 3.81 |
| **Gmet_1697** | *fabZ-1* | 4.26 | 5.85 | 4.97 | 4.20 |
| **Gmet_2059** | | 4.07 | 3.79 | 3.71 | 4.23 |
| **Gmet_1691** | *fabD-1* | 6.25 | 5.98 | 6.01 | 5.35 |
| **Gmet_1696** | | 7.34 | 8.37 | 8.43 | 7.07 |
| **Gmet_0287** | *fabB* | 4.80 | 5.03 | 5.21 | 7.27 |
| **Gmet_1694** | *fabG-1* | 7.69 | 8.21 | 7.94 | 7.44 |
| **Gmet_1613** | | 8.10 | 8.74 | 8.86 | 7.47 |
| **Gmet_1695** | *fabF-1* | 6.73 | 8.65 | 8.49 | 7.55 |
| **Gmet_2354** | *fabZ-2* | 7.98 | 8.21 | 8.29 | 7.67 |
| **Gmet_2558** | *fabI* | 8.08 | 8.00 | 7.86 | 7.67 |
| **Gmet_1216** | *accA* | 8.68 | 8.53 | 8.59 | 8.53 |
| **Gmet_1599** | *fabH-2* | 9.51 | 8.90 | 8.75 | 9.04 |
| **Gmet_3272** | *fabH-1* | 7.80 | 9.84 | 9.90 | 9.05 |
| **Gmet_1601** | *fabG-2* | 10.01 | 9.36 | 9.64 | 9.43 |
| **Gmet_2476** | *accD* | 8.66 | 9.46 | 8.89 | 9.64 |
| **Gmet_0984** | *accC* | 9.64 | 10.15 | 10.21 | 10.40 |
| **Gmet_1603** | *fabF-2* | 11.76 | 10.58 | 10.65 | 11.48 |
| **Gmet_0983** | *accB* | ND | ND | ND | ND |
| **Gmet_1600** | *fabD-2* | ND | ND | ND | ND |
| **Gmet_2229** | | ND | ND | ND | ND |
| **Gmet_2246** | | ND | ND | ND | ND |
| | Average | 7.65 | 8.00 | 7.91 | 7.63 |

Supplementary Table S3R. Average normalized TMM $\log_2$ values for *G. metallireducens* genes involved in glycolysis and gluconeogenesis

| Locus ID | Gene | GM/MB | GM/MA | GM/MS | GM/GS |
|---|---|---|---|---|---|

| | | | | | |
|---|---|---|---|---|---|
| **Gmet_2638** | *pckA* | 9.89 | 10.29 | 10.85 | 11.70 |
| **Gmet_3169** | | 7.80 | 8.14 | 8.12 | 6.55 |
| **Gmet_0122** | *pyk* | 8.09 | 9.21 | 8.83 | 10.25 |
| **Gmet_2372** | *eno* | 11.94 | 11.77 | 12.13 | 12.02 |
| **Gmet_1428** | | 8.74 | 8.45 | 8.44 | 8.98 |
| **Gmet_3203** | *gpmI* | 8.24 | 9.49 | 9.62 | 9.49 |
| **Gmet_1653** | *gpmA* | 7.48 | 7.05 | 7.08 | 4.92 |
| **Gmet_1947** | *pgk* | 8.10 | 8.33 | 8.22 | 9.47 |
| **Gmet_1211** | *gapB* | 7.82 | 8.41 | 8.30 | 7.53 |
| **Gmet_1946** | *gapA* | 10.39 | 9.88 | 10.55 | 11.58 |
| **Gmet_0789** | | 6.58 | 7.26 | 7.20 | 6.09 |
| **Gmet_1948** | *tpi* | ND | ND | ND | ND |
| **Gmet_1761** | | 9.05 | 8.80 | 9.06 | 9.62 |
| **Gmet_2383** | | 10.04 | 10.43 | 10.32 | 10.72 |
| **Gmet_1640** | *pfk-1* | 7.93 | 7.90 | 8.20 | 6.92 |
| **Gmet_0938** | *pfk-2* | 7.43 | 8.20 | 8.21 | 9.18 |
| **Gmet_1921** | *fbp* | 10.50 | 9.48 | 9.85 | 10.75 |
| **Gmet_2458** | *pgi* | 6.55 | 8.31 | 7.96 | 7.01 |
| **Gmet_1639** | | 8.03 | 7.60 | 8.21 | 6.48 |
| | Average | 8.59 | 8.83 | 8.95 | 8.85 |

Supplementary Table S3S. Average normalized TMM $\log_2$ values for *G. metallireducens* genes involved in the TCA cycle

| Locus ID | Gene | GM/MB | GM/MA | GM/MS | GM/GS |
|---|---|---|---|---|---|
| **Gmet_2752** | *aceF* | 7.71 | 9.33 | 9.18 | 7.73 |
| **Gmet_1016** | *acnA* | 7.31 | 9.50 | 8.54 | 10.70 |
| **Gmet_1912** | *acnB* | 11.18 | 11.86 | 11.72 | 10.92 |
| **Gmet_2340** | *acsA* | 6.85 | 7.25 | 7.79 | 6.11 |
| **Gmet_2509** | *bkdA* | 6.85 | 8.16 | 7.08 | 7.61 |
| **Gmet_2510** | *bkdB* | 6.42 | 8.46 | 7.59 | 6.63 |
| **Gmet_2511** | *bkdF* | 6.70 | 8.14 | 7.41 | 6.63 |
| **Gmet_2396** | *frdA* | 12.73 | 13.38 | 12.96 | 13.71 |
| **Gmet_2395** | *frdB* | 10.22 | 12.15 | 11.35 | 11.14 |
| **Gmet_2397** | *frdC* | 11.58 | 11.36 | 11.31 | 11.88 |
| **Gmet_2570** | *fumB* | 10.88 | 12.16 | 12.01 | 11.14 |
| **Gmet_2689** | *gltA* | 13.22 | 14.84 | 15.06 | 13.06 |

| | | | | | |
|---|---|---|---|---|---|
| **Gmet_1359** | *icd* | 13.83 | 13.52 | 13.58 | 13.15 |
| **Gmet_1362** | *korA* | 13.01 | 12.61 | 12.73 | 12.28 |
| **Gmet_1362** | *korA* | 13.01 | 12.61 | 12.73 | 12.28 |
| **Gmet_1363** | *korB* | 11.35 | 11.71 | 12.07 | 10.53 |
| **Gmet_1363** | *korB* | 11.35 | 11.71 | 12.07 | 10.53 |
| **Gmet_1364** | *korC* | 11.58 | 11.16 | 11.44 | 10.26 |
| **Gmet_1361** | *korD* | 10.36 | 8.37 | 9.24 | 11.28 |
| **Gmet_2764** | *lpdA-1* | 8.24 | 9.26 | 8.54 | 8.97 |
| **Gmet_1896** | *lpdA-2* | 5.94 | 5.13 | 4.40 | 5.54 |
| **Gmet_1360** | *mdh* | ND | ND | ND | ND |
| **Gmet_2760** | *pdhA* | 6.51 | 7.74 | 6.85 | 7.70 |
| **Gmet_2753** | *pdhB* | 7.62 | 8.02 | 7.57 | 9.69 |
| **Gmet_3419** | *por* | 10.25 | 10.76 | 11.29 | 13.34 |
| **Gmet_0816** | *pyc* | 11.64 | 11.27 | 11.39 | 11.67 |
| **Gmet_0308** | *sdhA* | 5.03 | 3.64 | 3.77 | 4.47 |
| **Gmet_0309** | *sdhB* | 4.91 | 3.46 | 3.83 | 4.11 |
| **Gmet_0310** | *sdhE* | 3.85 | 3.95 | 5.16 | 2.26 |
| **Gmet_2769** | *sucA* | 8.44 | 9.79 | 9.12 | 10.76 |
| **Gmet_2766** | *sucB* | 7.99 | 8.29 | 7.48 | 8.80 |
| **Gmet_0729** | *sucC-1* | 5.99 | 6.82 | 6.04 | 11.03 |
| **Gmet_2069** | *sucC-2* | 4.50 | 2.93 | 3.93 | 4.60 |
| **Gmet_2261** | *sucC-3* | ND | ND | ND | ND |
| **Gmet_0730** | *sucD-1* | 5.26 | 6.56 | 5.65 | 10.09 |
| **Gmet_2068** | *sucD-2* | 3.25 | 3.05 | 3.13 | 3.38 |
| **Gmet_2260** | *sucD-3* | ND | ND | ND | ND |
| **Gmet_1307** | *vorA* | 7.80 | 8.72 | 8.56 | 8.44 |
| **Gmet_1307** | *vorA* | 7.80 | 8.72 | 8.56 | 8.44 |
| **Gmet_1308** | *vorB* | 7.05 | 7.90 | 7.82 | 8.25 |
| **Gmet_1308** | *vorB* | 7.05 | 7.90 | 7.82 | 8.25 |
| **Gmet_1309** | *vorC* | 7.84 | 7.39 | 7.36 | 7.59 |
| **Gmet_1306** | *vorD* | 6.65 | 5.63 | 6.02 | 6.35 |
| **Gmet_1124** | | 4.63 | 5.63 | 5.42 | 6.19 |
| **Gmet_2763** | | 7.62 | 9.69 | 8.87 | 9.56 |
| | Average | 8.38 | 8.82 | 8.68 | 8.98 |

Supplementary Table S3T. Average normalized TMM $log_2$ values for *G. metallireducens* genes involved in oxidative phosphorylation

| Locus ID | Gene | GM/MB | GM/MA | GM/MS | GM/GS |
|---|---|---|---|---|---|
| **Gmet_0152** | *nuoA-2* | 5.07 | 7.39 | 6.58 | 6.43 |
| **Gmet_0153** | *nuoBCD* | 8.15 | 10.49 | 10.02 | 9.52 |
| **Gmet_0154** | *nuoE-2* | ND | ND | ND | ND |
| **Gmet_0157** | *nuoF-2* | 7.81 | 10.14 | 9.69 | 9.66 |
| **Gmet_0159** | *nuoG-2* | ND | ND | ND | ND |
| **Gmet_0161** | *nuoH-2* | 9.80 | 10.95 | 10.51 | 10.35 |
| **Gmet_0163** | *nuoI-2* | 6.62 | 7.33 | 7.09 | 7.87 |
| **Gmet_0165** | *nuoJ-2* | 5.40 | 6.94 | 6.93 | 7.00 |
| **Gmet_0167** | *nuoK-2* | 7.50 | 9.84 | 9.91 | 6.48 |
| **Gmet_0168** | *nuoL-2* | 8.52 | 9.91 | 9.42 | 8.44 |
| **Gmet_0171** | *nuoM-2* | 7.48 | 9.26 | 8.93 | 7.69 |
| **Gmet_0172** | *nuoN-2* | 7.03 | 8.97 | 8.92 | 8.59 |
| **Gmet_3342** | *nuoN-1* | 12.97 | 12.02 | 12.64 | 12.00 |
| **Gmet_3343** | *nuoM-1* | 12.52 | 11.96 | 12.35 | 11.69 |
| **Gmet_3344** | *nuoL-1* | 12.47 | 12.35 | 12.79 | 11.81 |
| **Gmet_3345** | *nuoK-1* | 10.64 | 8.50 | 9.61 | 10.10 |
| **Gmet_3346** | *nuoJ-1* | 11.70 | 10.39 | 10.83 | 11.55 |
| **Gmet_3347** | *nuoI-1* | 12.37 | 9.69 | 10.67 | 11.29 |
| **Gmet_3348** | *nuoH-1* | 11.49 | 11.37 | 11.78 | 10.76 |
| **Gmet_3349** | *nuoG-1* | ND | ND | ND | ND |
| **Gmet_3350** | *nuoF-1* | 12.78 | 13.01 | 13.42 | 12.20 |
| **Gmet_3351** | *nuoE-1* | 12.09 | 11.24 | 11.79 | 12.19 |
| **Gmet_3352** | *nuoD* | 12.48 | 12.63 | 12.95 | 12.60 |
| **Gmet_3353** | *nuoC* | 12.11 | 11.00 | 11.54 | 12.05 |
| **Gmet_3354** | *nuoB* | 10.64 | 11.54 | 11.83 | 11.30 |
| **Gmet_3355** | *nuoA-1* | 12.64 | 10.37 | 11.44 | 11.13 |
| **Gmet_3359** | *atpB* | 9.66 | 9.44 | 9.54 | 10.92 |
| **Gmet_3360** | *atpE* | 10.30 | 10.55 | 10.34 | 10.85 |
| **Gmet_3405** | *atpC* | 9.92 | 9.22 | 9.27 | 10.97 |
| **Gmet_3406** | *atpD* | 10.44 | 11.01 | 10.94 | 12.78 |
| **Gmet_3407** | *atpG* | 9.39 | 9.62 | 9.76 | 10.30 |
| **Gmet_3408** | *atpA* | 10.63 | 11.22 | 11.24 | 12.32 |

| | | | | | |
|---|---|---|---|---|---|
| **Gmet_3409** | *atpH* | 9.98 | 9.17 | 9.32 | 11.71 |
| **Gmet_3410** | *atpF* | ND | ND | ND | ND |
| **Gmet_3411** | *atpX* | 9.84 | 8.94 | 9.19 | 9.45 |
| **Gmet_3497** | | ND | ND | ND | ND |
| | Average | 10.01 | 10.21 | 10.36 | 10.39 |

Supplementary Table S3U. Average normalized TMM $\log_2$ values for *G. metallireducens* genes involved in purine biosynthesis

| Locus ID | Gene | GM/MB | GM/MA | GM/MS | GM/GS |
|---|---|---|---|---|---|
| **Gmet_2848** | *prsA* | 9.54 | 9.59 | 9.98 | 9.32 |
| **Gmet_0048** | *nudF* | 6.95 | 8.24 | 8.32 | 6.80 |
| **Gmet_1943** | *purB* | 7.84 | 8.92 | 8.67 | 8.78 |
| **Gmet_3260** | *purA* | 10.12 | 10.49 | 10.68 | 10.07 |
| **Gmet_1181** | | 8.63 | 8.00 | 7.69 | 8.42 |
| **Gmet_0647** | *adk* | 12.01 | 10.84 | 11.03 | 11.24 |
| **Gmet_2686** | *ndk* | 10.71 | 9.61 | 9.68 | 11.05 |
| **Gmet_1297** | *nrdJ* | 8.26 | 8.66 | 8.95 | 9.36 |
| **Gmet_1910** | | 6.03 | 5.59 | 5.84 | 3.91 |
| **Gmet_1638** | | 8.20 | 6.15 | 6.99 | 5.24 |
| **Gmet_2859** | *cysD* | 5.55 | 6.22 | 5.03 | 8.87 |
| **Gmet_2293** | *guaB* | 8.81 | 9.73 | 9.59 | 10.99 |
| **Gmet_2292** | *guaA* | 8.65 | 9.53 | 9.53 | 11.04 |
| **Gmet_1630** | | 5.14 | 4.93 | 5.27 | 5.34 |
| **Gmet_1778** | | 5.32 | 6.50 | 5.97 | 6.38 |
| **Gmet_3434** | | 6.43 | 8.16 | 8.59 | 8.79 |
| **Gmet_2716** | | 4.90 | 5.90 | 5.92 | 4.50 |
| **Gmet_2327** | *gmk* | 9.25 | 9.20 | 9.39 | 10.71 |
| **Gmet_1762** | | 8.51 | 7.76 | 7.87 | 7.43 |
| **Gmet_1939** | *purF* | 8.08 | 8.47 | 8.53 | 9.33 |
| **Gmet_2904** | *purD* | 7.12 | 8.68 | 8.52 | 6.71 |
| **Gmet_1845** | *purN* | 9.28 | 8.03 | 7.83 | 8.23 |
| **Gmet_3193** | *purT* | 7.60 | 8.94 | 8.87 | 8.19 |
| **Gmet_1940** | *purQ* | 6.94 | 8.30 | 8.19 | 8.88 |
| **Gmet_1941** | *purSL* | 9.79 | 10.47 | 10.20 | 10.64 |
| **Gmet_1844** | *purM* | 9.05 | 9.36 | 9.16 | 7.69 |

| | | | | | |
|---|---|---|---|---|---|
| **Gmet_2903** | *purE* | 5.96 | 6.96 | 6.43 | 5.83 |
| **Gmet_0916** | *purC* | 9.03 | 8.88 | 9.19 | 10.89 |
| **Gmet_1422** | *apt* | 9.35 | 7.64 | 7.85 | 8.69 |
| **Gmet_2905** | *purH* | 7.49 | 8.87 | 8.49 | 8.43 |
| **Gmet_1419** | *surE* | 5.79 | 5.87 | 5.91 | 5.86 |
| **Gmet_1778** | | 5.32 | 6.50 | 5.97 | 6.38 |
| **Gmet_3434** | | 6.43 | 8.16 | 8.59 | 8.79 |
| **Gmet_2716** | | 4.90 | 5.90 | 5.92 | 4.50 |
| **Gmet_2201** | | ND | ND | ND | ND |
| **Gmet_2219** | | ND | ND | ND | ND |
| **Gmet_1875** | *rdgB* | 6.17 | 7.08 | 6.51 | 5.88 |
| | Average | 7.69 | 8.06 | 8.03 | 8.09 |

Supplementary Table S3V. Average normalized TMM $\log_2$ values for *G. metallireducens* genes involved in pyrimidine biosynthesis

| Locus ID | Gene | GM/MB | GM/MA | GM/MS | GM/GS |
|---|---|---|---|---|---|
| **Gmet_0661** | *carB-2* | 8.57 | 9.60 | 9.53 | 8.11 |
| **Gmet_1771** | *carA* | 8.95 | 8.90 | 8.88 | 9.78 |
| **Gmet_1774** | *carB-1* | 9.85 | 9.85 | 9.91 | 10.96 |
| **Gmet_1769** | *pyrB* | 8.65 | 8.83 | 8.77 | 8.84 |
| **Gmet_1770** | *pyrC* | 9.49 | 9.23 | 9.36 | 9.56 |
| **Gmet_1841** | *pyrD* | 7.58 | 7.57 | 7.97 | 7.79 |
| **Gmet_1842** | *pyrK* | 7.23 | 7.48 | 7.94 | 6.78 |
| **Gmet_1938** | *pyrE* | 7.35 | 7.27 | 7.20 | 8.89 |
| **Gmet_1355** | *pyrF* | 8.86 | 7.94 | 7.63 | 7.66 |
| **Gmet_1252** | *pyrH* | 9.79 | 8.95 | 9.01 | 9.77 |
| **Gmet_2686** | *ndk* | 10.71 | 9.61 | 9.68 | 11.05 |
| **Gmet_1276** | *pyrG* | 9.43 | 9.91 | 9.80 | 9.33 |
| **Gmet_0895** | *maf* | 6.70 | 7.33 | 7.59 | 8.14 |
| **Gmet_0865** | *cmk* | 7.19 | 7.71 | 7.91 | 6.81 |
| **Gmet_1768** | *pyrR* | 8.49 | 7.93 | 8.38 | 9.04 |
| **Gmet_1778** | | ND | ND | ND | ND |
| **Gmet_1181** | | 8.63 | 8.00 | 7.69 | 8.42 |
| **Gmet_1297** | *nrdJ* | 8.26 | 8.66 | 8.95 | 9.36 |
| **Gmet_1910** | | 6.03 | 5.59 | 5.84 | 3.91 |

| | | | | | |
|---|---|---|---|---|---|
| **Gmet_1622** | | 5.93 | 7.16 | 7.11 | 7.88 |
| **Gmet_2318** | *tmk-1* | ND | ND | ND | ND |
| **Gmet_3250** | *tmk-2* | 6.99 | 7.82 | 8.34 | 8.90 |
| **Gmet_0378** | *thyX* | 9.08 | 9.66 | 10.08 | 10.09 |
| | Average | 8.27 | 8.33 | 8.45 | 8.62 |

Supplementary Table S4. Fold difference between the transcripts for pilin and pilin accessory genes and the median $log_2$ TMM values for normalized *G. metallireducens* transcripts from RNAseq libraries assembled from co-cultures with *G. metallireducens* and *G. sulfurreducens* (GM/GS), *G. metallireducens* and *M. barkeri* (GM/MB), *G. metallireducens* and *M. acetivorans* (GM/MA), and *G. metallireducens* and *M. subterranea* (GM/MS). Standard deviations were calculated from triplicate samples. NS: no significant difference.

| Locus ID | Gene | GM/GS | GM/MB | GM/MA | GM/MS |
|---|---|---|---|---|---|
| **Gmet_0734** | *pilV-1* | 2.05 +/- 0.08 | 1.98 +/- 0.006 | 2.05 +/- 0.08 | 2.14 +/- 0.12 |
| **Gmet_0735** | *pilW-1* | 2.06+/-0.09 | 2.07 +/- 0.05 | 2.34 +/- 0.06 | 2.47 +/- 0.09 |
| **Gmet_0736** | *pilX-1* | 1.90 +/- 0.005 | 1.95 +/- 0.03 | 2.10 +/- 0.05 | 2.09 +/- 0.07 |
| **Gmet_0737** | *pilY-1* | 2.41 +/- 0.04 | 2.45 +/- 0.03 | 2.73 +/- 0.14 | 2.87 +/- 0.09 |
| **Gmet_0959** | *pilD* | 1.82 +/- 0.07 | 1.37 +/- 0.38 | 2.11 +/- 0.11 | 2.20 +/- 0.10 |
| **Gmet_0962** | *pilE* | 2.55 +/- 0.05 | NS | NS | NS |
| **Gmet_0964** | *pilV-2* | 2.19 +/- 0.02 | NS | NS | NS |
| **Gmet_0965** | *pilW-2* | 2.29 +/- 0.06 | NS | NS | NS |
| **Gmet_0966** | *pilX-2* | 2.39 +/- 0.05 | NS | NS | NS |
| **Gmet_0967** | *pilY-2* | 2.94 +/- 0.02 | 2.43 +/- 0.03 | 2.18 +/- 0.06 | 2.00 +/- 0.06 |
| **Gmet_0971** | *pilM* | 2.57 +/- 0.06 | 2.43 +/- 0.06 | 2.51 +/- 0.12 | 2.43 +/- 0.15 |
| **Gmet_0972** | *pilN* | 2.15 +/- 0.02 | 2.21 +/- 0.05 | 2.22 +/- 0.10 | 2.11 +/- 0.08 |
| **Gmet_0973** | *pilO* | 2.48 +/- 0.06 | 2.31 +/- 0.06 | 2.22 +/- 0.09 | 2.14 +/- 0.07 |
| **Gmet_0974** | *pilP* | 2.03 +/- 0.06 | 2.11 +/- 0.04 | 2.16 +/- 0.10 | 2.19 +/- 0.11 |
| **Gmet_0975** | *pilQ* | 2.94 +/- 0.04 | 2.99 +/- 0.09 | 2.87 +/- 0.12 | 2.86 +/- 0.08 |
| **Gmet_1393** | *pilB* | 2.66+/- 0.03 | 2.38 +/- 0.05 | 2.18 +/- 0.06 | 2.11 +/- 0.03 |
| **Gmet_1394** | *pilT-4* | 2.56 +/- 0.03 | 2.28 +/- 0.07 | 2.09 +/- 0.07 | 2.05 +/- 0.06 |
| **Gmet_1395** | *pilC* | 2.53 +/- 0.06 | 2.32 +/- 0.04 | 2.17 +/- 0.08 | 2.08 +/- 0.06 |
| **Gmet_1396** | *pilS* | 2.25 +/- 0.02 | 2.32 +/- 0.06 | 2.08 +/- 0.06 | 2.05 +/- 0.01 |
| **Gmet_1397** | *pilR* | 2.18 +/- 0.02 | 2.33 +/- 0.04 | 2.08 +/- 0.09 | 2.06 +/- 0.05 |
| **Gmet_1399** | *pilA* | 2.34 +/- 0.13 | 2.90 +/- 0.10 | 2.15 +/- 0.06 | 2.14 +/- 0.09 |
| **Gmet_1400** | *spc* | 2.24 +/- 0.05 | 3.01 +/- 0.09 | 2.11 +/- 0.20 | 2.22 +/- 0.12 |

Supplementary Table S5A. Summary of statistics from RNAseq libraries assembled from RNA extracted from co-cultures of *M. acetivorans* and *G. metallireducens* cells grown with ethanol (20 mM) provided as the electron donor for DIET (3 biological replicates).

| | Mean | SEM | Max | Min | Units |
|---|---|---|---|---|---|
| **Raw (unfiltered) reads** | 84,371,021 | 2,983,587 | 92,482,194 | 76,141,695 | #reads/library |
| **QC-filtered merged paired end reads** | 47,870,723 | 2,656,797 | 51,201,184 | 42,619,863 | #reads/library |
| **QC-filtered merged paired mRNA reads** | 6,614,197 | 587,282 | 7,728,580 | 5,735,580 | #mRNA reads/library |
| **QC-filtered merged paired end read size** | 154.37 | 1.05 | 190 | 100 | Bases/read |

Supplementary Table S5B. Summary of statistics from RNAseq libraries assembled from RNA extracted from co-cultures of *M. subterranea* and *G. metallireducens* cells grown with ethanol (20 mM) provided as the electron donor for DIET (3 biological replicates).

| | Mean | SEM | Max | Min | Units |
|---|---|---|---|---|---|
| **Raw (unfiltered) reads** | 81,582,527 | 2,797,437 | 85,561,268 | 76,186,665 | #reads/library |
| **QC-filtered merged paired end reads** | 80,039,493 | 2,581,745 | 83,825,440 | 75,105,767 | #reads/library |
| **QC-filtered merged paired mRNA reads** | 7,625,132 | 744,989 | 8,711,430 | 6,198,810 | #mRNA reads/library |
| **QC-filtered merged paired end read size** | 128 | 15.7 | 175 | 90 | Bases/read |

Supplementary Table S5C. Summary of statistics from RNAseq libraries assembled from RNA extracted from co-cultures of *M. barkeri* and *G. metallireducens* cells grown with ethanol (20 mM) provided as the electron donor for DIET (3 biological replicates).

| | Mean | SEM | Max | Min | Units |
|---|---|---|---|---|---|
| **Raw (unfiltered) reads** | 40,617,676 | 1,910,602 | 44,430,082 | 38,487,036 | #reads/library |
| **QC-filtered merged paired end reads** | 23,088,289 | 1,396,820 | 25,836,238 | 21,278,931 | #reads/library |
| **QC-filtered merged paired mRNA reads** | 7,513,666 | 262,469 | 8,029,516 | 7,171,584 | #mRNA reads/library |
| **QC-filtered merged paired end read size** | 145 | 36.8 | 190 | 100 | Bases/read |

Supplementary Table S5D. Summary of statistics from RNAseq libraries assembled from RNA extracted from co-cultures of *G. sulfurreducens* and *G. metallireducens* cells grown with ethanol (10 mM) provided as the electron donor and fumarate (45 mM) provided as the electron acceptor for DIET (3 biological replicates).

| | Mean | SEM | Max | Min | Units |
|---|---|---|---|---|---|
| **Raw (unfiltered) reads** | 13,175,156 | 718,065 | 15,400,547 | 10,989,034 | #reads/library |
| **QC-filtered merged paired end reads** | 9,286,241 | 961,335 | 10,951,323 | 7,621,159 | #reads/library |
| **QC-filtered merged paired mRNA reads** | 1,219,374 | 701,931 | 1,473,813 | 1,035,632 | #mRNA reads/library |
| **QC-filtered merged paired end read size** | 106.9 | 4.72 | 135 | 100 | Bases/read |

Supplementary Table S6. Fold differences for genes coding for flagellin and flagellum assembly proteins in *G. metallireducens* in co-cultures grown with *G. sulfurreducens* (GS) compared to co-cultures grown with the 3 different *Methanosarcina* species (*M. barkeri* (MB); *M. acetivorans* (MA); *M. subterranea* (MS)). p-values are available in Supplementary Table S2 but all comparisons shown have p-values that are <0.05. NS: no significant difference

| Locus ID | Annotation | Gene | GS/MB | GS/MA | GS/MS |
|---|---|---|---|---|---|
| **Gmet_0426** | Flagellar biosynthesis protein FlhA | *flhA* | 2.00 | 2.10 | 2.82 |
| **Gmet_0427** | Flagellar biosynthesis protein FlhF | *flhF* | NS | NS | 2.05 |
| **Gmet_0428** | Flagellar biosynthesis protein FlhG | *flhG* | 1.58 | NS | 1.72 |
| **Gmet_0429** | RNA polymerase sigma factor FliA | *fliA* | NS | NS | 2.13 |
| **Gmet_0430** | Flagellar basal-body rod protein FlgF | *flgF* | 2.01 | 2.11 | 2.81 |
| **Gmet_0431** | Flagellar basal-body rod protein FlgG | *flgG* | 1.46 | 1.44 | 2.26 |
| **Gmet_0432** | Flagellar basal body P-ring formation protein FlgA | *flgA* | 2.18 | NS | NS |
| **Gmet_0433** | Flagellar L-ring protein FlgH | *flgH* | NS | NS | NS |
| **Gmet_0434** | Flagellar P-ring protein precursor | *flgI* | 2.09 | NS | 1.55 |
| **Gmet_0435** | Peptidoglycan hydrolase FlgJ | *flgJ* | NS | 3.16 | 4.88 |
| **Gmet_0436** | Anti-sigma-28 factor, FlgM | *flgM* | 3.63 | 7.89 | 14.53 |
| **Gmet_0437** | FlgN-like domain | *flgN* | 1.41 | 2.17 | 3.88 |
| **Gmet_0438** | Flagellar hook-associated protein | *flgK* | 1.60 | 3.77 | 6.44 |
| **Gmet_0439** | Flagellar hook-associated protein 3 FlgL | *flgL* | 1.51 | 4.26 | 4.95 |
| **Gmet_0440** | Carbon storage regulator, CsrA | *csrA* | 2.10 | 5.33 | 6.99 |
| **Gmet_0441** | Flagellar assembly factor FliW | *fliW* | NS | NS | 1.50 |
| **Gmet_0442** | Flagellin protein | *fliC* | 12.81 | 31.21 | 33.43 |
| **Gmet_0444** | Flagellar hook-associated protein 2 | *fliD* | 1.58 | 2.45 | 1.48 |
| **Gmet_0445** | Flagellar secretion chaperone FliS | *fliS* | 2.32 | 5.80 | 3.04 |
| **Gmet_0446** | Flagellar protein FliT | *fliT* | 2.19 | 5.26 | 3.55 |
